# Controlled administration of aerosolized SARS-CoV-2 to K18-hACE2 transgenic mice uncouples respiratory infection and anosmia from fatal neuroinvasion

**DOI:** 10.1101/2021.08.06.455382

**Authors:** Valeria Fumagalli, Micol Ravà, Davide Marotta, Pietro Di Lucia, Chiara Laura, Eleonora Sala, Marta Grillo, Elisa Bono, Leonardo Giustini, Chiara Perucchini, Marta Mainetti, Alessandro Sessa, José M. Garcia-Manteiga, Lorena Donnici, Lara Manganaro, Serena Delbue, Vania Broccoli, Raffaele De Francesco, Patrizia D’Adamo, Mirela Kuka, Luca G. Guidotti, Matteo Iannacone

## Abstract

The development of a tractable small animal model faithfully reproducing human COVID-19 pathogenesis would arguably meet a pressing need in biomedical research. Thus far, most investigators have used transgenic mice expressing the human ACE2 in epithelial cells (K18-hACE2 transgenic mice) that are intranasally instilled with a liquid SARS-CoV-2 suspension under deep anesthesia. Unfortunately, this experimental approach results in disproportionate high CNS infection leading to fatal encephalitis, which is rarely observed in humans and severely limits this model’s usefulness. Here, we describe the use of an inhalation tower system that allows exposure of unanesthetized mice to aerosolized virus under controlled conditions. Aerosol exposure of K18-hACE2 transgenic mice to SARS-CoV-2 resulted in robust viral replication in the respiratory tract, anosmia, and airway obstruction, but did not lead to fatal viral neuroinvasion. When compared to intranasal inoculation, aerosol infection resulted in a more pronounced lung pathology including increased immune infiltration, fibrin deposition and a transcriptional signature comparable to that observed in SARS-CoV-2- infected patients. This model may prove useful for studies of viral transmission, disease pathogenesis (including long-term consequences of SARS-CoV-2 infection) and therapeutic interventions.

## Main text

The coronavirus disease 2019 (COVID-19) pandemic is caused by the recently identified *β*-coronavirus severe acute respiratory syndrome coronavirus 2 (SARS-CoV-2) (Wu et al., 2020; Zhou et al., 2020). Disease severity is variable, ranging from asymptomatic infection to multi-organ failure and death. Though SARS-CoV-2 primarily targets the respiratory system, some patients with COVID-19 can also exhibit extrarespiratory symptoms, including neurological manifestations such as loss of smell (anosmia) and taste (ageusia), headache, fatigue, memory impairment, vomiting, gait disorders and impaired consciousness (Helms et al., 2020; Qiu et al., 2020; Xydakis et al., 2020; Iadecola et al., 2020). SARS-CoV-2 can infect neurons in human brain organoids (Ramani et al., 2020; Song et al., 2021) and a few studies reported the presence of SARS-CoV-2 in olfactory sensory neurons (OSNs) and deeper areas within the central nervous system (CNS) in fatal COVID-19 cases (Glatzel et al., 2021; Matschke et al., 2020; Puelles et al., 2020; Song et al., 2021; Melo et al., 2021). However, the neurotropism of SARS-CoV-2 and a direct role of CNS infection in the pathogenesis of neurological manifestations remains highly debated.

Despite the availability of effective vaccines against SARS-CoV-2, we still know little about COVID-19 pathogenesis. Arguably, the availability of tractable animal models to mechanistically dissect virological, immunological and pathogenetic aspects of the infection with SARS-CoV-2 and future human coronaviruses would be a major benefit.

Wild-type laboratory mice are poorly susceptible to SARS-CoV-2 infection because the mouse angiotensin-converting enzyme (ACE) 2 does not act as a cellular receptor for the virus (Muñoz-Fontela et al., 2020). Several transgenic mouse lineages expressing the human version of the SARS-CoV-2 receptor (hACE2) support viral replication and recapitulate certain clinical characteristics of the human infection (Muñoz-Fontela et al., 2020). The most widely used model is the K18-hACE2 transgenic mouse (McCray et al., 2007), which expresses hACE2 predominantly in epithelial cells under the control of the cytokeratin 18 (*KRT18*) promoter (Chow et al., 1997). K18-hACE2 mice are typically infected by intranasally instilling liquid suspensions of SARS-CoV-2 under deep anesthesia. This results in disproportionate high CNS infection leading to fatal encephalitis (Jiang et al., 2020; Sun et al., 2020; Winkler et al., 2020; Zheng et al., 2021; Kumari et al., 2021), which rarely occurs in patients with COVID-19. Such viral neuroinvasion severely limits the usefulness of these mouse models, hampering studies on disease pathogenesis (including long-term consequences of SARS-CoV-2 infection) as well as on drug discovery.

SARS-CoV-2 is mainly transmitted from person to person via respiratory droplets (Zhou et al., 2021). In an attempt to mimic this transmission route, we made use of a nose-only inhalation tower system that allows to expose unanesthetized mice to aerosolized virus under controlled pressure, temperature and humidity conditions (see **Figure S1A-C** and Methods). Animals were located inside a restraint with a neck clip positioned between the base of the skull and the shoulders, thus avoiding thorax compression, keeping the airways completely unobstructed and allowing for spontaneous breathing through the nose. K18-hACE2 transgenic mice were infected with a target dose of 1 × 10^5^ tissue culture infectious dose 50 (TCID50) of SARS-CoV-2 either through intranasal administration with 25 μl of diluted virus (IN) or through a 20 to 30 minute exposure to aerosolized virus (AR) (**Figure 1A-B**, see Methods). Pulmonary function was measured during aerosol exposure using plethysmography. Frequency, tidal volume, minute volume and accumulated volume of SARS-CoV-2-exposed mice were comparable to PBS-exposed mice (**Figure S1D**).

**Figure 1.**
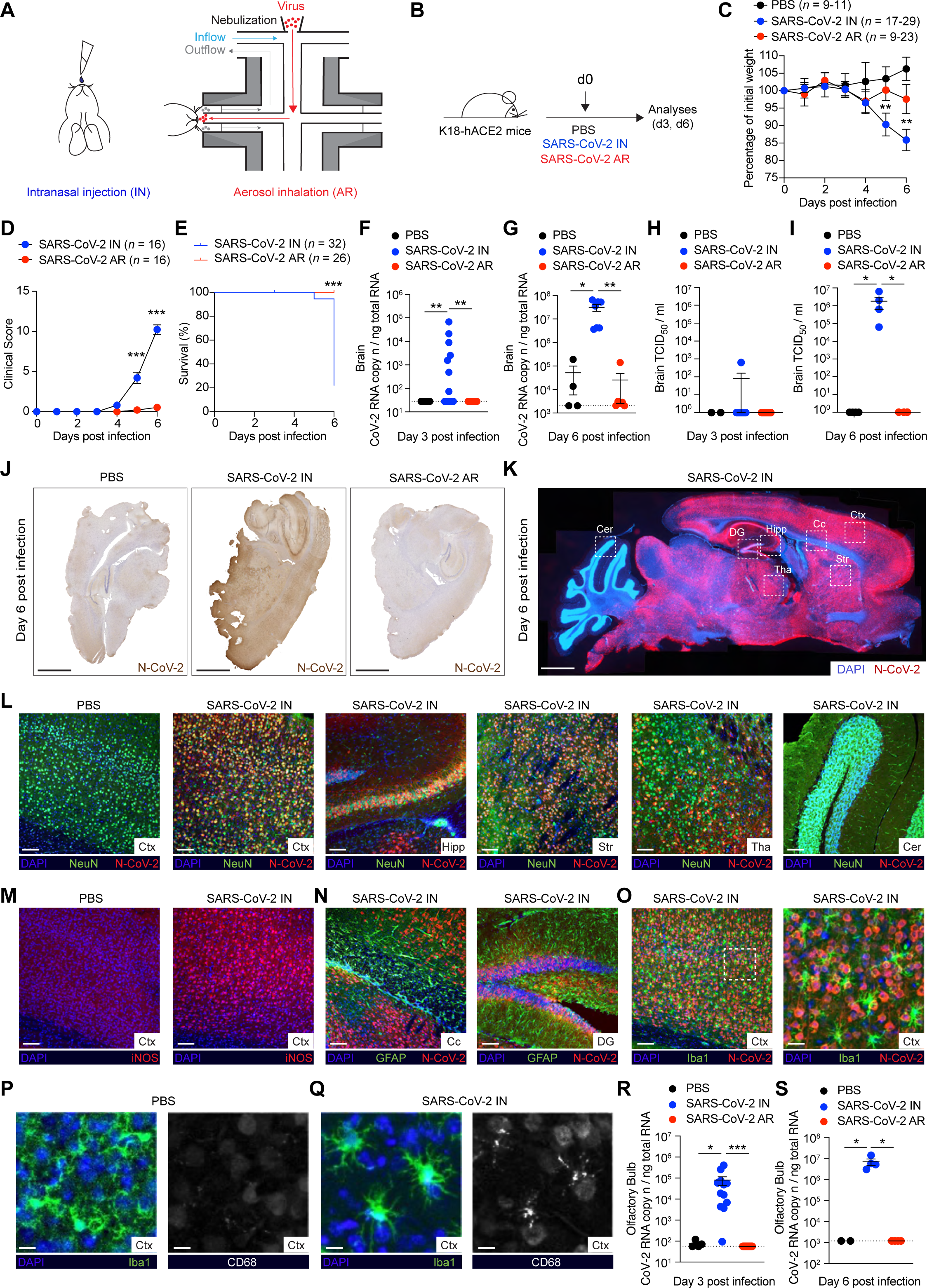
Intranasal inoculation, but not aerosol exposure, of SARS-CoV-2 leads to fatal neuroinvasion in K18-hACE2 transgenic mice. (A) Illustration of the two modalities used to infect K18-hACE2 mice with SARS-CoV-2. On the left, intranasal injection (IN) is shown. On the right, representation of an unanesthetized mouse placed in the nose-only Allay restrainer on the inhalation chamber is shown. In red, aerosolized virus with a particle size of ∼4 μm; in light-blue, primary flow set to 0,5 L/min/port; in grey, mouse breathing outflow (see Methods). **(B)** Schematic representation of the experimental setup. K18-hACE2 mice were infected with a target dose of 1 x 10^5^ TCID50 of SARS-CoV-2 through intranasal (IN) administration or through aerosol (AR) exposure. Lung, brain, nasal turbinates, olfactory bulbs, and blood were collected and analyzed 3 days and 6 days post infection. **(C)** Mouse body weight was monitored daily for up to 6 days and is expressed as the percentage of weight relative to the initial weight on day 0. Statistical significance of comparison between IN- (*n* = 17-29, blue dots) and AR-infected mice (*n* = 9-23, red dots) is shown. Control mice treated with PBS are also shown (*n* = 9-11, black dots). Data are represented as mean ± SEM. **(D)** Clinical score was assessed evaluating the piloerection (0-3), posture (0-3), activity level (0-3), eye closure (0-3) and breathing (0-3) (see Methods). Statistical significance of comparison between IN- (*n* = 16, blue dots) and AR-infected mice (*n* = 16, red dots) is shown. PBS-treated control mice did not exhibit any clinical signs. **(E)** Survival curve of IN- (*n* = 32, blue dots) and AR-infected mice (*n* = 26, red dots). **(F, G)** Quantification of SARS-CoV-2 RNA in the brain of IN- (*n* = 7-11, blue dots) and AR- (*n* = 6-10, red dots) infected mice as well as of PBS-treated control mice (*n* = 4, black dots) measured 3 days (**F**) and 6 days (**G**) post infection. RNA values are expressed as copy number per ng of total RNA and the limit of detection is indicated as a dotted line. **(H, I)** Viral titers in the brain were determined 3 (**H**) and 6 days (**I**) after infection by median tissue culture infectious dose (TCID50). PBS-treated control mice: *n* = 2-3, black dots; IN- infected mice: *n* = 4, blue dots; AR-infected mice: *n* = 4, red dots. **(J)** Representative immunohistochemical micrographs of sagittal brain sections from PBS-treated control mice (left), IN- (middle) and AR-infected mice (right) 6 days post infection. N-SARS- CoV-2 positive cells are depicted in brown. Scale bars, 1 mm. **(K)** Representative confocal immunofluorescence staining for N-CoV-2 (red) in sagittal brain sections of IN- infected mice. Cell nuclei are depicted in blue. White boxes indicate different brain areas: cerebellum (Cer); dentate gyrus (DG); hippocampus (Hipp); corpus callosum (Cc); cerebral cortex (Ctx); thalamus (Tha); striatum (Str). Scale bars, 1 mm. **(L)** Representative confocal immunofluorescence micrographs of sagittal brain sections from PBS-treated control mice (first panel) and IN-infected mice 6 days post infection. N-CoV-2 is depicted in red, NeuN neural marker in green and cell nuclei in blue. Fields of cerebral cortex (Ctx), hippocampus (Hipp), striatum (Str), thalamus (Tha) and cerebellum (Cer) are shown. Scale bars, 100 μm. **(M)** Representative confocal immunofluorescence micrographs of cerebral cortex (Ctx) in PBS-treated control mice (left panel) and IN-infected mice (right panel) 6 days post infection. iNOS^+^ cells are depicted in red and cell nuclei in blue. Scale bars, 100 μm. **(N)** Representative confocal immunofluorescence micrographs of two areas of the brain from IN-infected mice 6 days post infection. Corpus callosum (Cc) of the cerebral cortex, left panel, and dentate gyrus (DG), right panel. N-CoV-2 is depicted in red, GFAP astroglial marker in green and cell nuclei in blue. Scale bars, 100 μm. **(O)** Representative confocal immunofluorescence micrographs of the cerebral cortex (Ctx) of IN-infected mice at 6 days post infection. N-CoV-2 is depicted in red, Iba1 microglial marker in green and cell nuclei in blue. White box indicates the magnification represented in the right panel. Scale bars represent 100 μm (image) and 25 μm (magnification). **(P, Q)** Representative confocal immunofluorescence micrographs of the cerebral cortex (Ctx) of PBS-treated control mice (**P**) and IN-infected mice (**Q**) 6 days post infection. Left panels show Iba1 microglial marker in green and cell nuclei in blue; right panel shows CD68 marker of microglial activation in white. Scale bars, 10 μm. **(R, S)** Quantification of SARS-CoV-2 RNA in the olfactory bulbs of IN- (*n* = 4-12, blue dots) and AR- (*n* = 4-12, red dots) infected mice as well as of PBS-treated control mice (*n* = 2-4, black dots) measured 3 days (**R**) and 6 days (**S**) post infection. RNA values are expressed as copy number per ng of total RNA and the limit of detection is indicated as a dotted line. Data are expressed as mean ± SEM. Data in (C-I, R, S) are pooled from 2 independent experiments per time point. * p-value < 0.05, ** p-value < 0.01, *** p-value < 0.001; two- way ANOVA followed by Sidak’s multiple comparison test (**C, D,** comparison between blue and red dots for each time point); Log-rank (Mantel-Cox) test (**E**); Kruskal-Wallis test (**F**-**I, R-S**).

As expected (Zheng et al., 2021; Kumari et al., 2021), IN-infected animals exhibited significant body weight loss and a severe clinical score (see Methods for details), so that, by day 6 post infection (p.i.), ∼ 80% of them had died and the remaining ones appeared lethargic (**Figure 1C-E**). By contrast, AR-infected mice maintained stable body weight, and did not show any signs of disease nor mortality (**Figure 1C-E**). The severe disease observed in IN-infected K18-hACE2 transgenic mice was associated with the detection of high viral RNA titers and infectious virus in the brain (**Figure 1F- I**). By contrast, neither SARS-CoV-2 RNA nor infectious virus were detected in the brain of mice exposed to aerosolized virus (**Figure 1F-I**). Immunohistochemical and immunofluorescence staining confirmed the presence of the SARS-CoV-2 nucleoprotein in the brain of IN-infected, but not AR-infected, mice (**Figure 1J, K** and data not shown). Specifically, diffuse staining for SARS-CoV-2 nucleoprotein was detected throughout the cerebrum with comparable staining in the different brain areas with the notable exception of the cerebellum where most of its cells stained negative for viral antigens (**Figure 1K**). Neurons were by far the most infected brain cells as shown by the co-staining of the SARS-CoV-2 nucleoprotein with the pan-neuronal marker NeuN (*∼*90% double positive cells, **Figure 1L** and **Figure S2A**). Accordingly, neurons in infected brains strongly up-regulated iNOS (Goody et al., 2005), which was undetectable in neuronal cells from control, uninfected mice (**Figure 1M**). By contrast, only a minor fraction of astrocytes (*∼*2%) and microglia (*∼*4%) stained positive for the SARS-CoV-2 nucleoprotein (**Figure 1N, O** and **Figure S2B, C**). Of note, Iba1^+^ myeloid cells in SARS-CoV-2-infected brains were activated as revealed by the characteristic morphology (swollen processes with reduced ramifications) and CD68 positivity (**Figure 1P, Q**). Consistent with the data on the recovery of infectious virus, viral RNA, and viral antigens, we found a significant immune cell recruitment (particularly of T cells, B cells, monocytes, and eosinophils) in the brains of IN-infected, but not AR-infected mice (**Figure S2D, E**). Together, these results reveal a profound viral neuroinvasion which correlates with the severe health deterioration in IN-infected mice. The high viral load and widespread viral distribution in the brain of IN-infected mice contrasts with the occasional localized detection of SARS-CoV-2 in the olfactory bulbs and/or the medulla of fatal COVID-19 cases (Bradley et al., 2020; Kantonen et al., 2020; Matschke et al., 2020; Meinhardt et al., 2021; Reichard et al., 2020; Serrano et al., 2021) and caution against utilizing this model to investigate the neurological complications of SARS-CoV-2 infection in humans.

We next investigated the potential SARS-CoV-2 entry portals to the central nervous system (CNS) in IN-infected K18-hACE2 transgenic mice. One possibility is that the virus gains access to the CNS via the blood-brain barrier, which implies a viremic phase. However, no SARS-CoV-2 RNA was ever detected in the sera of infected mice (**Figure S2F, G**), consistent with earlier reports (Winkler et al., 2020; Zheng et al., 2021; Kumari et al., 2021). Alternatively, SARS-CoV-2 could enter the CNS by retrograde axonal transport upon olfactory sensory neuron infection. Indeed, and in line with previous studies (Winkler et al., 2020; Zheng et al., 2021; Kumari et al., 2021), viral RNA was detected in the olfactory bulb of IN-infected, but not AR-infected, mice (**Figure 1R, S**). Overall, the data indicate that IN, but not AR, infection of K18- hACE2 transgenic mice with SARS-CoV-2 results in lethal neuroinvasion likely via retrograde axonal transport after olfactory sensory neuron infection.

We next analyzed viral replication in the upper respiratory tract of K18-hACE2 transgenic mice infected with SARS-CoV-2 via intranasal inoculation or aerosol exposure. We detected the presence of SARS-CoV-2 RNA in the nasal turbinates of both AR- and IN-infected mice at days 3 and 6 p.i. (**Figure 2A, B**). In order to examine whether viral replication within the upper respiratory tract induced anosmia, we subjected AR- and IN-infected mice to a social scent-discrimination assay (Zheng et al., 2021) (**Figure 2C**). If olfaction is normal (as in PBS-treated controls), male mice exposed to tubes containing male or female bedding preferentially spend time sniffing the female scent (**Figure 2D-F**). By contrast, both AR- as well as IN-infected mice spent significantly less time sniffing the female scent at day 3 p.i. (**Figure 2D, E**), indicative of hyposmia or anosmia. Importantly, at day 3 p.i. mobility of both AR- and IN-infected mice was normal, as there were no differences in the amount of time spent sniffing the male tube (**Figure 2D**). At day 6 p.i., AR-infected mice still showed signs of hyposmia/anosmia, whereas IN-infected mice were completely lethargic preventing further analyses (**Figure 2D-F**). The data obtained in AR-infected mice are consistent with the hypothesis that hyposmia or anosmia occur because of the infection of olfactory epithelium and in the absence of CNS infection or general malaise (Zheng et al., 2021).

**Figure 2.**
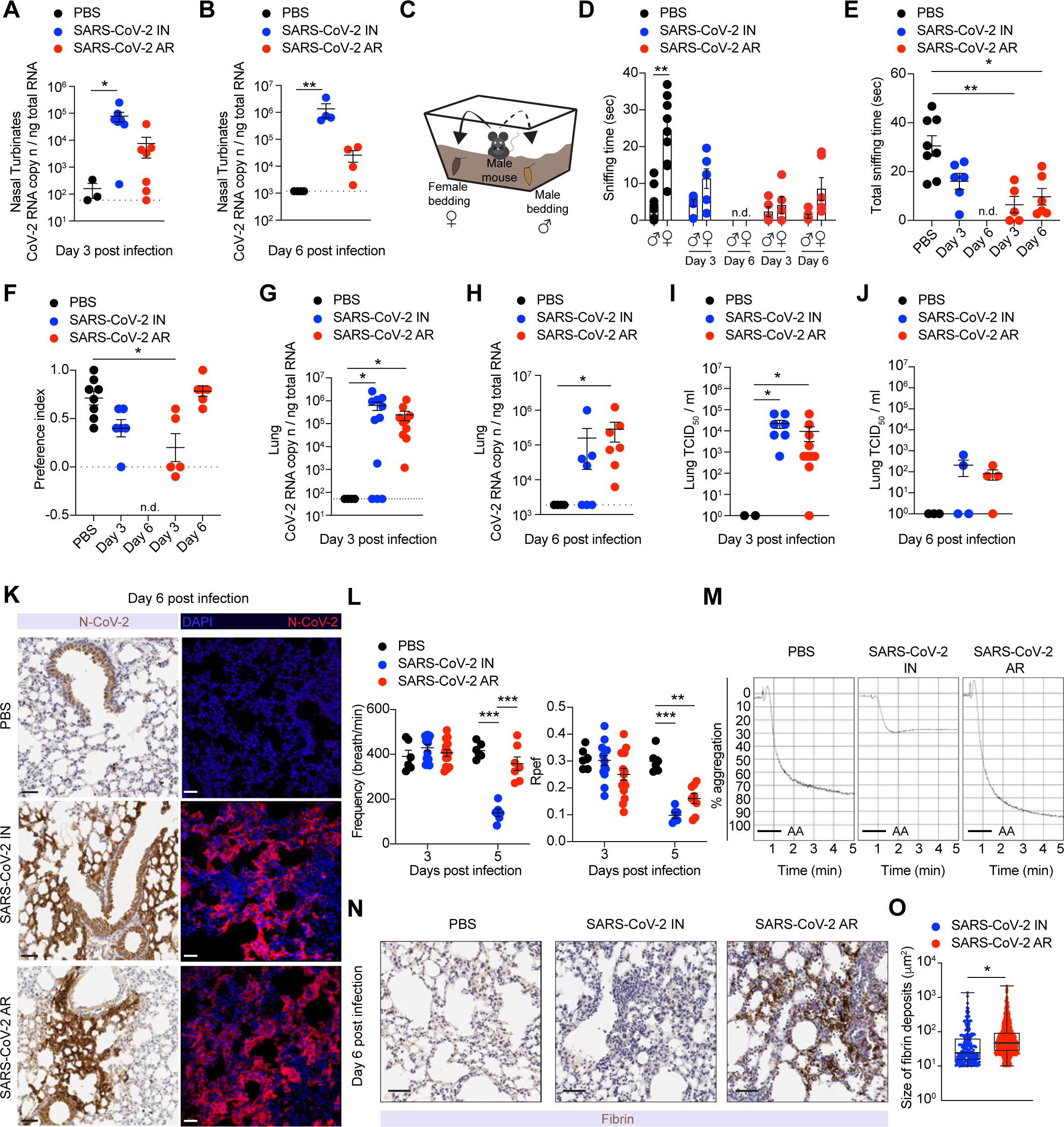
Aerosol exposure of K18-hACE2 transgenic mice to SARS-CoV-2 leads to efficient respiratory infection, anosmia, and fibrin deposition in the lung. (A-B) Quantification of SARS-CoV-2 RNA in the nasal turbinates of PBS-treated control mice (*n* = 3, black dots) and of IN- (*n* = 7, blue dots) and AR- (*n* = 7, red dots) infected mice 3 days (**A**) and 6 days (**B**) post infection. RNA values are expressed as copy number per ng of total RNA and the limit of detection is indicated as a dotted line. **(C)** Illustration showing social scent-discrimination test. Male mice were free to investigate for 5 minutes two different tubes containing their own cage bedding or female cage bedding placed at two opposite corners of a clean cage. **(D, E)** Time that males spent sniffing their own male scent or female scent (**D**) and the sum of the two times (**E**) is expressed as time sniffing (seconds). Analyses were performed 3 or 6 days post IN- (*n* = 3-6, blue dots) or AR-infection (*n* = 3-5, red dots). As control, PBS-treated mice are shown (*n* = 8, black dots). In D, comparison between female and male time in each group of mice. n.d, the sniffing time could not be determined since mice were completely lethargic. **(F)** Preference index for male mice was calculated as (female time – male time)/ (total sniffing time female + male), as an indicator of the time spent sniffing preferred (female) or non-preferred (male) scents. n.d.: the sniffing time could not be determined since mice were completely lethargic. **(G, H)** Quantification of SARS-CoV-2 RNA in the lung of IN- (*n* = 7-11, blue dots) and AR- (*n* = 7-10, red dots) infected mice as well as ofPBS-treated control mice (*n* = 4, black dots) measured 3 days (**G**) and 6 days (**H**) post infection. RNA values are expressed as copy number per ng of total RNA and the limit of detection is indicated as a dotted line. **(I, J)** Viral titers in the lung were determined 3 (**I**) and 6 days (**J**) after infection by median tissue culture infectious dose (TCID50). PBS-treated control mice: *n* = 2-3, black dots; IN-infected mice: *n* = 4-7, blue dots; AR- infected mice: *n* = 4-10, red dots. **(K)** Representative immunohistochemical (left) and confocal immunofluorescence (right) micrographs of lung sections from PBS-treated control mice (top), IN- (middle) and AR-infected mice (bottom) at 6 days post infection. N-CoV-2 positive cells are depicted in brown (left panels) or in red (right panels). Cell nuclei are depicted in blue (right panels). Scale bars, 30 μm. **(L)** Pulmonary function was assessed by whole-body plethysmography performed 3 and 5 days post IN- (*n* = 6- 14, blue dots) and AR-infection (*n* = 7-14, red dots). As control, PBS-treated mice were evaluated (*n* = 6, black dots). Frequency (left) and Rpef (right) parameters are shown. Calculated respiratory values were averaged over a 15 minute-data collection period. **(M)** Representative aggregometry curves induced by arachidonic acid (AA) on platelet- rich plasma from PBS-treated control mice (left), IN- (middle) and AR-infected mice (right) 6 days post infection. Platelet aggregation was measured by light transmission aggregometry for 5 minutes and is expressed as % aggregation. **(N)** Representative immunohistochemical micrographs of lung sections from PBS-treated control mice (left), IN- (middle) and AR-infected mice (right) at 6 days post infection. Fibrin deposition is shown in brown. Scale bars, 30 μm. **(O)** Quantification of the size of fibrin deposits (μm^2^). *n* = 4. Data are expressed as mean ± SEM and are pooled from 2 independent experiments per time point. * p-value < 0.05, ** p-value < 0.01, *** p-value < 0.001; Kruskal-Wallis test (**A, B, E**-**J**); two-way ANOVA followed by Sidak’s multiple comparison test (**L**); Mann-Whitney U-test two-tailed (**D, O**). In **D**, statistical analysis was performed comparing female and male sniffing time within the same experimental group of mice.

We next assessed viral replication in the lower respiratory tract of SARS-CoV-2- infected K18-hACE2 transgenic mice. We detected comparable amounts of SARS-CoV- 2 RNA and infectious virus from the lungs of mice infected with the two different routes of administration at both day 3 and day 6 p.i. (**Figure 2G-J**). Immunohistochemical and immunofluorescence staining for the SARS-CoV-2 nucleoprotein confirmed similar levels of viral antigens and similar staining patterns in the lungs of IN- and AR-infected mice (**Figure 2K**). To gain insight into the impact of infection on lung physiology, pulmonary function was measured at day 3 and 5 p.i. via whole-body plethysmography. Consistent with previously published data (Leist et al., 2020; Menachery et al., 2015), we confirmed that, when compared to control mice, IN-infected mice exhibited a significant loss in pulmonary function as indicated by changes in e.g., respiratory frequency, tidal volume, Rpef (a measure of airway obstruction) and PenH (a controversial metric that has been used by some as an indirect measure of airway resistance and by others as a non-specific assessment of breathing patterns (Bates et al., 2004; Lundblad et al., 2007; Lomask, 2006; Menachery et al., 2015)) (**Figure 2L** and **Figure S3A**). Interestingly, most of these respiratory parameters were normal or much less altered in AR-infected mice, suggesting that the observed changes in IN- infected mice were mostly due to CNS infection. One notable exception is Rpef, a calculated index of airway resistance that considers the time needed to reach maximum expiratory flow and the total expiratory time (Menachery et al., 2015). Interestingly, Rpef was the only metric that was consistently altered in AR-infected mice (and to the same extent as in IN-infected mice), suggesting that this index might truly reflect lung infection and pathology, rather than CNS involvement.

COVID-19, particularly in its most severe forms, has been associated with thrombotic phenomena that entail increased platelet activation and aggregation, and fibrin deposition in the lungs (Lax et al., 2020; Fox et al., 2020; Dolhnikoff et al., 2020; Mast et al., 2021; Manne et al., 2020). We therefore set out to assess platelet function by performing light transmission aggregometry of platelet rich plasma (PRP) obtained from infected mice. Whereas IN infection resulted in a significantly impaired platelet aggregation, PRP from AR-infected mice showed a normal or even increased aggregation (**Figure 2M**). Interestingly, this was associated with increased fibrin deposition and larger platelet aggregates in the lungs of AR-infected mice (**Figure 2N, O** and **Figure S3B**). Together, the data indicate that aerosol exposure of K18-hACE2 transgenic mice to SARS-CoV-2 results in robust viral replication in the respiratory tract, anosmia, airway obstruction, and platelet aggregation with fibrin deposition in the lung.

Although the vigorous SARS-CoV-2 replication in the lungs of AR-infected K18-hACE2 transgenic mice suggests that these mice *de facto* inhaled at least the same amount of virus as IN-infected mice, it is theoretically possible that a higher viral inoculum would have resulted in fatal neuroinvasion even upon aerosol delivery. However, when we infected mice with 5 × 10^5^ TCID50 of SARS-CoV-2 via aerosol delivery, we still failed to detect any viral replication in the CNS (data not shown). To further increase viral replication in the infected hosts, we transiently inhibited type 1 IFN receptor signaling with anti-IFNAR1 blocking antibodies (Abs) prior to AR infection (**Figure S4A**). As expected, higher levels of viral RNA (**Figure S4B**) and infectious virus (**Figure S4C**) were detected in the lungs of mice treated with anti-IFNAR1 Abs compared to control mice. The presence of higher amounts of virus in the lungs of anti-IFNAR1-treated mice was confirmed by immunohistochemical staining for the SARS-CoV-2 nucleoprotein (**Figure S4D**). Despite the increased lung viral load, we failed to detect SARS-CoV-2 RNA, infectious virus, or viral antigens in the brain of anti-IFNAR1-treated AR-infected mice (**Figure S4E-G**).

We next sought to better characterize the histopathological changes and the immune response in the lungs of SARS-CoV-2 infected mice. Hematoxylin and eosin staining revealed an inflammatory process that peaked at day 6 post infection and appeared more severe in AR- than in IN-infected mice (**Figure 3A** and **Figure S5A**). Lung sections revealed interstitial edema, consolidation, alveolar wall thickening and immune infiltration of both polymorphonuclear and mononuclear cells in the alveolar as well as the interstitial space (**Figure 3A** and **Figure S5A**). Consistent with the histology, the absolute number of cells recovered 6 days after infection from the lung and bronchoalveolar lavage (BAL) was significantly higher in AR-infected than in IN-infected mice (**Figure 3B-E**). Specifically, the differences in immune cell recruitment could be imputed to an increase in CD4^+^ and CD8^+^ T cells as well as in monocytes (**Figure 3B- E**). A higher number of TCR-*β*^+^ T cells in the lung of AR-infected mice was confirmed by confocal immunofluorescence histology (**Figure S5B**).

**Figure 3.**
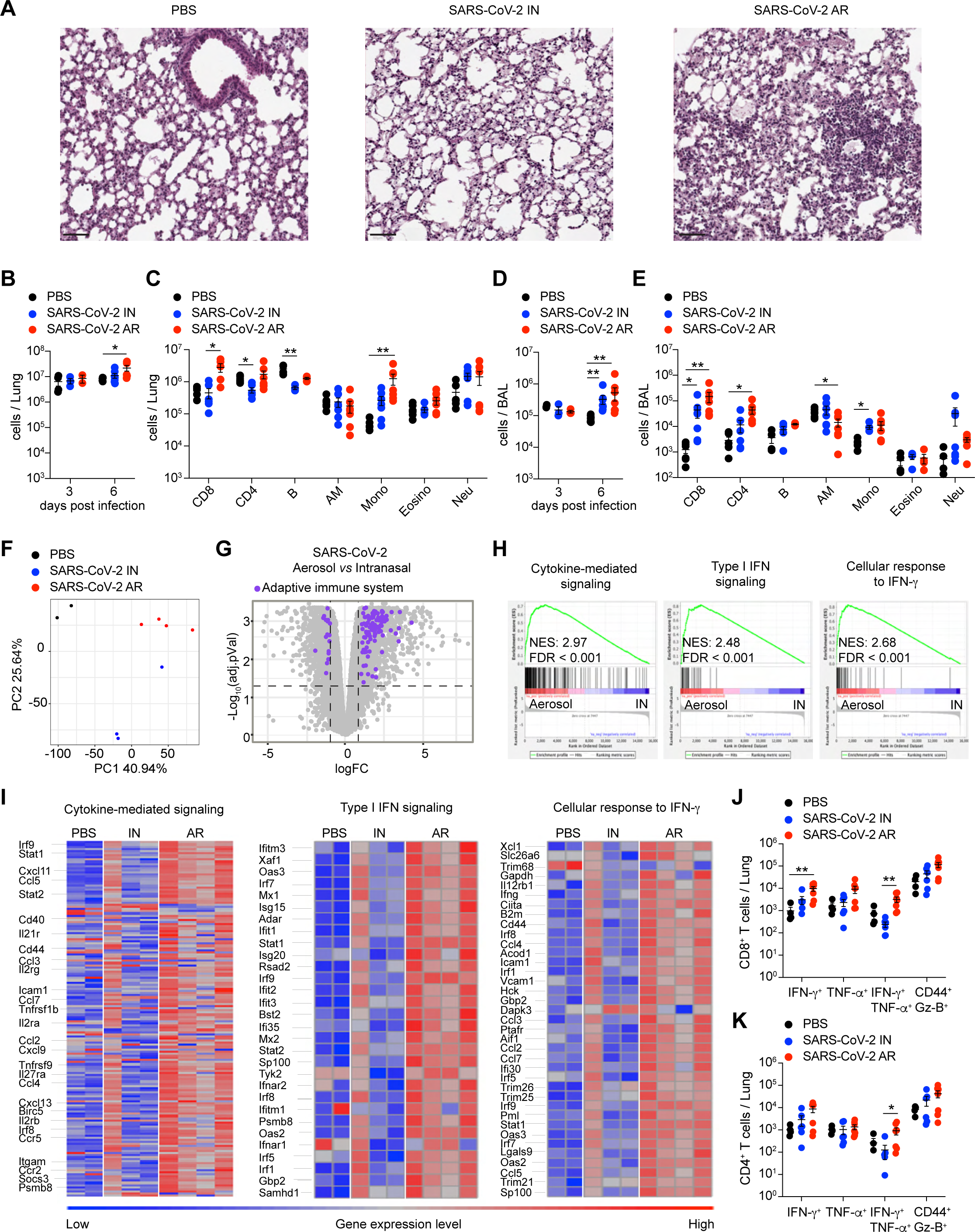
Histopathological changes, immune response, and transcriptional signatures in the lungs of infected mice. **(A)** Representative hematoxylin/eosin (H&E) micrographs of lung sections from PBS-treated control mice (left), IN- (middle) and AR-infected mice (right) 6 days post infection. Scale bars, 50 μm. **(B-E)** Absolute number of total cells (**B, D**) and of single cell populations (**C, E**) recovered from lung homogenates (**B, C**) and bronchoalveolar lavage (BAL) (**D, E**) of PBS-treated control mice (*n* = 4-6, black dots), IN- (*n* = 3-7, blue dots) and AR-infected mice (*n* = 3-7, red dots) analyzed 6 days post infection. CD8^+^ T cells (Live, CD45^+^, CD8^+^); CD4^+^ T cells (Live, CD45^+^, CD4^+^); B cells (Live, CD45^+^, CD8^-^, CD4^-^, B220^+^, CD19^+^); AM, alveolar macrophages (Live, CD45^+^, CD8^-^, CD4^-^, Ly6g^-^, CD11b^-^, F4/80^+^, SiglecF^hi^); Mono, monocytes (Live, CD45^+^, CD8^-^, CD4^-^, Ly6g^-^, SiglecF^-^, CD11b^+^, Ly-6c^+^); Eosino, eosinophils (Live, CD45^+^, CD8^-^, CD4^-^, Ly6g^-^, CD11b^+^, SiglecF^int^); Neu, neutrophils (Live, CD45^+^, CD8^-^, CD4^-^, CD11b^+^, Ly6g^+^). **(F)** Principal Component Analysis (PCA) of RNA-seq expression values from the lungs of PBS-treated, control (*n* = 2, black), IN- (*n* = 3, blue) and AR-infected (*n* = 4, red) mice. Percentages indicate the variance explained by each PC. **(G)** Volcano plot of RNA-Seq results. The X-axis represents the Log2 Fold-Change of Differentially Expressed Genes (DEG) comparing AR- to IN-infection, the Y-axis the -Log10(FDR). Genes significantly upregulated in AR- relative to IN- (|logFC| > 1 and adjusted P value < 0.05, horizontal and vertical dashed line) belonging to the “Adaptive immune system” pathway from BioPlanet 2019 (Huang et al., 2019) are highlighted in violet. **(H)** GSEA of three gene sets described in (Winkler et al., 2020), comparing the transcriptome of IN- and AR-infected mice, presented as the running enrichment score for the gene set as the analysis ’walks down’ the ranked list of genes (reflective of the degree to which the gene set is over-represented at the top or bottom of the ranked list of genes) (top), the position of the gene-set members (black vertical lines) in the ranked list of genes (middle) and the value of the ranking metric (bottom). **(I)** Heatmaps of genes (one per row) belonging to the three signatures as in (**H**), expressed in logarithmic normalized read counts. Each column represents an individual sample and results were visualized using the pheatmap R package. **(J, K)** Absolute number of CD8^+^ (**J**) and CD4^+^ (**K**) T cells producing IFN-*γ*, TNF-*α* or both and expressing CD44 and Granzyme-B (Gz-B) in the lung of PBS-treated control mice (*n* = 4, black dots), IN- (*n* = 5, blue dots) and AR-infected mice (*n* = 5, red dots) 6 days after infection. Data are expressed as mean ± SEM. Data in (B-E, J, K) are pooled from 2 independent experiments. * p-value < 0.05, ** p-value < 0.01; Kruskal-Wallis test (**B-E, J, K**).

Next, we sought to analyze the transcriptome in the lung of infected K18-hACE2 mice by performing bulk RNA sequencing (RNA-seq) of lung homogenates 6 days after SARS-CoV-2 infection. Principal component analysis revealed distinct transcriptional signatures between AR-infected, IN-infected, and uninfected mice (**Figure 3F**). Genes upregulated upon AR infection were associated to adaptive immune responses, and to immune system signaling by interferons and other cytokines (**Figure 3G**, **Figure S6** and **Table S1**). In particular, the transcription of genes related to cytokine-mediated signaling, type I IFN signaling and cellular response to IFN-*γ* (Winkler et al., 2020) was increased in the lung of AR-infected mice with respect to IN-infected ones (**Figure 3H, I** and **Table S1**). Interestingly, many human orthologs of the genes upregulated in the lungs of AR-infected mice were found to be also induced in COVID-19 patients, including genes related to leukocyte trafficking (e.g., *Ccl11*, *Ccl8*, *Ccl2*, *Cxcl9* and *Cxcl10*), antiviral response induced by type I IFN (*Ifitm3*, *Ifit2*, *Oas1a*, *Oas3*, *Stat1*, *Irf1*, *Mx1*, *Mx2*, *Isg15*) and TNF (*Tnfsf10*) (Delorey et al., 2021; Blanco-Melo et al., 2020; Karki et al., 2021)(**Figure 3I**, **Figure S6** and **Table S1**). Of note, *Ccl8* and *Ccl2* as well as *Cxcl9* and *Cxcl10*, chemoattractants for monocytes and T cells, respectively, were significantly upregulated in the lungs of AR-infected mice and in COVID-19 patients, in line with the increased recruitment of these cells (**Figure 3I**, **Figure S6C** and **Table S1**). The enrichment of the gene signature related to the cellular response to IFN-*γ* in AR- infected mice prompted us to assess SARS-CoV-2-specific T cell responses. Antigen- specific CD8^+^ and CD4^+^ T cells recovered from lung homogenates were assessed for IFN-*γ*, TNF-*α* and Granzyme-B (Gz-B) expression upon *in vitro* stimulation with the H2- D^b^-restricted S538-546 and I-A^b^-restricted ORF3a 266-280 immunodominant peptides (Zhuang et al., 2021). In line with the RNA-seq data, we found that the absolute number of IFN-*γ*^+^ TNF-*α*^+^ virus-specific CD8^+^ and CD4^+^ T cells were significantly higher in the lung of AR-infected mice compared to IN-infected mice (**Figure 3J, K**).

In summary, we have generated and characterized a novel COVID-19 platform, based on controlled aerosol exposure of K18-hACE2 transgenic mice to SARS-CoV-2. Mice infected via aerosol develop robust respiratory infection, anosmia, and signs of airway obstruction but, in contrast to mice infected intranasally, do not experience fatal neuroinvasion. Moreover, when compared to intranasal inoculation, aerosol exposure results in a more severe lung pathology, inflammation, and fibrin deposition. We believe that this model may allow studies on viral transmission (e.g., by analyzing the effect of aerosol particle size, humidity, and temperature on infectivity), on disease pathogenesis (including, potentially, thrombotic events and long-term consequences of infection) and on therapeutic interventions.

## Acknowledgments

We thank M. Freschi, M. Raso, A. Fiocchi, M. Genua, R. Ostuni for technical support; M. Silva for secretarial assistance; and the members of the Iannacone laboratory for helpful discussions. Flow cytometry was carried out at FRACTAL, a flow cytometry resource and advanced cytometry technical applications laboratory established by the San Raffaele Scientific Institute. Confocal immunofluorescence histology was carried out at Alembic, an advanced microscopy laboratory established by the San Raffaele Scientific Institute and the Vita-Salute San Raffaele University. We would like to acknowledge the PhD program in Basic and Applied Immunology and Oncology at Vita-Salute San Raffaele University, as D.M. and E.S. conducted this study as partial fulfillment of their PhD in Molecular Medicine within that program. M.I. is supported by the European Research Council (ERC) Consolidator Grant 725038, ERC Proof of Concept Grant 957502, Italian Association for Cancer Research (AIRC) Grants 19891 and 22737, Italian Ministry of Health Grants RF-2018-12365801 and COVID- 2020-12371617, Lombardy Foundation for Biomedical Research (FRRB) Grant 2015- 0010, the European Molecular Biology Organization Young Investigator Program, and Funded Research Agreements from Gilead Sciences, GSK Vaccines, Takis Biotech and Toscana Life Sciences. L.G.G. is supported by the Italian Association for Cancer Research (AIRC) Grant 22737, Lombardy Open Innovation Grant 229452, PRIN Grant 2017MPCWPY from the Italian Ministry of Education, University and Research, Funded Research Agreements from Gilead Sciences, Avalia Therapeutics and CNCCS SCARL and donations from FONDAZIONE SAME and FONDAZIONE PROSSIMO MIO for COVID-19-related research. M.K. is supported by the Italian Ministry of Education, University and Research grants SIR-RBSI14BAO5 and PRIN-2017ZXT5WR.

## Author contributions

Conceptualization, V.F., L.G.G., M.I.; Investigation, V.F., M.R., D.M., P.D.L., C.L., E.S., M.G., E.B., L.G., C.P., M.M., A.S., J.M.G-M.; Resources, L.D., L.M., S.D., R.D.F.; Formal Analysis, V.F., M.R., D.M., E.S., V.B., P.D’A., M.K.; Writing V.F., M.K., L.G.G., M.I. with input from all authors; Visualization, V.F.; Project supervision, M.I.; Funding Acquisition, L.G.G. and M.I.

## Data and materials availability

Accession numbers will be made available prior to publication.

## Materials and Methods

### Mice

B6.Cg-Tg(K18-ACE2)^2Prlmn/^J mice (referred to in the text as K18-hACE2) were purchased from The Jackson Laboratory. Mice were housed under specific pathogen-free conditions and heterozygous mice were used at 8-10 weeks of age. All experimental animal procedures were approved by the Institutional Animal Committee of the San Raffaele Scientific Institute and all infectious work was performed in designed BSL-3 workspaces.

### Virus

The SARS-CoV-2/human/ITA/Milan-UNIMI-1/2020 (GenBank: MT748758.1) isolation was carried out in BSL-3 workspace and performed in Vero E6 cells, which were cultured at 37°C, 5% CO2 in complete medium (DMEM supplemented with 10% FBS, MEM non-essential amino acids, 100 U/ml penicillin, 100 U/ml streptomycin, 2mM L-glutamine). Virus stocks were titrated using Endpoint Dilutions Assay (EDA, TCID50/ml). Vero E6 cells were seeded into 96 wells plates and infected at 95% of confluency with base 10 dilutions of virus stock. After 1h of adsorption at 37°C, the cell-free virus was removed, cells were washed with PBS 1X, and complete medium was added to cells. After 48h, plates were evaluated for the presence of a cytopathic effect (CPE). TCID50/ml of viral stocks were then determined by applying the Reed–Muench formula.

### Nose-only inhalation tower system

The DSI nose-only inhalation tower system (DSI Buxco respiratory solutions, DSI) is composed of 7 open ports where mice are exposed to aerosolized virus. Mice were placed in the DSI Allay restraint. The Allay collar is positioned between the base of the mouse skull and its shoulders, thus avoiding thorax compression, and maintaining normal breathing patterns. Liquid virus was aerosolized by an Aeroneb (Aerogen) vibrating mesh nebulizer that generates particles of ∼4 μm size which were uniformly delivered to all the tower ports. One port of the tower was occupied by a temperature and humidity probe for real-time monitoring of the tower conditions. The inhalation tower controller software was used to define the flow and pressure of the inhalation tower as well as the temperature, humidity, and the ratio O2/CO2 inside the tower. During virus exposure mice were monitored through plethysmography for frequency, tidal volume, minute volume and accumulated volume. The whole system was placed inside a class II biological safety hood that is located in a BSL-3 facility.

### Mouse infection through aerosol exposure (AR) or intranasal administration (IN)

Unanesthetized K18-hACE2 mice were placed in a nose-only Allay restrainer on the inhalation chamber (DSI Buxco respiratory solutions, DSI). To reach a target accumulated inhaled aerosol (also known as delivered dose) of 1 x 10^5^ TCID50, mice were exposed to aerosolized SARS-CoV-2 for 20-30 minutes (depending on the total volume of diluted virus and on the number of mice simultaneously exposed). Primary inflows and pressure were controlled and set to 0,5 L/minute/port and -0,5 cmH2O, respectively. As control, K18-hACE2 mice received the same volume of aerosolized PBS (125 μL per mouse).

Intranasal administration of 1 x 10^5^ TCID50 of SARS-COV-2 per mouse in a total volume of 25 μL PBS was performed under 2% isoflurane (#IsoVet250) anesthesia.

Infected mice were monitored daily to record body weight, clinical and respiratory parameters. The clinical score was based on a cumulative 0-3 scale evaluating fur, posture, activity level, eyes and breathing (see table below).

| Parameter | Degree | Scoring points |
| --- | --- | --- |
| Fur | Shining | 0 |
|  | Matte | 1 |
|  | Ruffled | 3 |
| Posture | Normal | 0 |
|  | Hunched | 1 |
|  | Massively hunched | 3 |
| Activity level | Active (social contacts) | 0 |
|  | Impaired (motility after stimulation) | 1 |
|  | No activity (lethargy) | 3 |
| Eyes | Clear and clean | 0 |
|  | Semi-closed | 1 |
|  | Unclean and closed | 3 |
| Breathing | Normal | 0 |
|  | Slightly changed | 1 |
|  | Strongly accelerated | 3 |

### *In vivo* treatment

In selected experiments, K18-hACE2 mice were injected intraperitoneally with 2 mg per mouse of *α*-IFNAR1 blocking antibody (BioXcell, #BE0241, clone MAR1-5A3) 1 day before infection.

### Social scent-discrimination test

The social scent-discrimination test was used to assess hyposmia/anosmia in IN-infected or AR-infected male mice compared to PBS-treated control male mice, as previously described (Zheng et al., 2021). Briefly, two 2 ml-Eppendorf tubes containing beddings from the cage of grouped females and from the experimental male were placed at two opposing corners of a clean cage. Experimental male mice were scored for 5 minutes for the time spent in sniffing the tubes, considering only the time when the nose was inside one of the two tubes. Control male mice preferentially explored the tube containing the female bedding. Preference index was calculated as: (time spent to sniff female tube - time spent to sniff male tube) / (time spent to sniff female tube + time spent to sniff male tube).

### Whole-body plethysmography

Whole-body plethysmography (WBP) was performed using WBP chamber (DSI Buxco respiratory solutions, DSI). Mice were allowed to acclimate inside the chamber for 10 minutes before recording respiratory parameters for 15 minutes using the FinePointe software.

### Platelet aggregation

Blood was collected from the retro-orbital sinus into 1:10 volume of citrate phosphate dextrose (CPD; Sigma-Aldrich, #C7165) and platelet-rich plasma (PRP) was prepared as described (Iannacone et al., 2008). The PRP platelet count was adjusted to the lowest value of the day. Homologous platelet-poor plasma (PPP) was isolated by spinning the remaining peripheral blood from PRP at 3500 rpm for 10 minutes.

Aggregation in stirred PRP at 37°C was induced by adding 125 μM of Arachidonic Acid (Mascia Brunelli, #311501WB) and monitored by recording changes in light transmittance through the PRP suspension using a Chrono-log model 490 aggregometer (Chrono-log Corporation, Havertown, PA).

### Cell Isolation and Flow Cytometry

Mice were anesthetized and the trachea was cannulated with a 22-G canula (BD, #381223) followed by bronchoalveolar lavage (BAL) with 3 washes of 1 ml sterile PBS. BAL fluid was centrifuged to obtain a single cell suspension. Mice were euthanized by cervical dislocation. At the time of autopsy, mice were perfused through the right ventricle with PBS. Brain and olfactory bulbs were removed from the skull and nasal turbinates from the nose cavity. Lung tissue was digested in RPMI 1640 containing 3.2 mg/ml Collagenase IV (Sigma, #C5138) and 25 U/ml DNAse I (Sigma, #D4263) for 30 minutes at 37°C. Brain was digested in RPMI 1640 containing 1 mg/ml Collagenase D (Sigma, # 11088866001) and 50 U/ml DNAse I for 30 minutes at 37°C. Homogenized lungs and brains were passed through 70 μm nylon meshes to obtain a single cell suspension. Cells were resuspended in 36% percoll solution (Sigma #P4937) and centrifuged for 20 minutes at 2000 rpm (light acceleration and low brake). The remaining red blood cells were removed with ACK lysis.

For analysis of *ex-vivo* intracellular cytokine production, 1 mg/ml of brefeldin A (Sigma #B7651) was included in the digestion buffer. All flow cytometry stainings of surface-expressed and intracellular molecules were performed as described (Bénéchet et al., 2019). Briefly, cells were stimulated for 4h at 37°C in the presence of brefeldin A, monensin (life technologies, #00-4505-51) and two immunodominant peptides covering the ORF3a 266-280 (restricted to I-A^b^, EPIYDEPTTTTSVPL) and the spike protein S538-546 (restricted to H2-D^b^, CVNFNFNGL) of SARS-CoV-2, as described (Zhuang et al., 2021). Cell viability was assessed by staining with Viobility™ 405/520 fixable dye (Miltenyi, Cat #130-109-814). Antibodies (Abs) are indicated in the table below.

| Name | Clone | Source and catalog number |
| --- | --- | --- |
| CD103 | 2E7 | Biolegend #121408 |
| CD62L | RM4-5 | Biolegend #104420, Biolegend #104428 |
| CD11b | M1/70 | Biolegend #101218 |
| IA/IE | M5/114.15.2 | Biolegend #107622 |
| Ly-6c | HK1.4 | Biolegend #128013 |
| Siglec-F | E50-2440 | BD Biosciences #740388 |
| CD64 | X54-5/7.1 | BD OptiBuild #740622 |
| CD8 | 53-6.7 | Biolegend #100725, Biolegend #100759 |
| CD11c | HL3 | BD Biosciences #563735 |
| CD4 | RM4-5 | BD Biosciences #740208; BD Biosciences #741912 |
| B220 | RA3-6B2 | BD Biosciences #564662 |
| CD44 | IM7 | BD Biosciences #741227; BioLegend #103028 |
| CD69 | H1.2F3 | BD Biosciences #612793 |
| CD19 | 1D3 | BD Biosciences #749027 |
| F4/80 | BM8 | Biolegend #123110 |
| Ly-6g | 1A5 | BD Pharmingen #562700 |
| CD45 | 30-F11 | Biolegend #103113, BD Biosciences #564279 |
| IFN-γ | XMG1.2 | Biolegend #505813 |
| Bcl-6 | K112-91 | Biolegend #581525 |
| TNF-α | MP6-XT22 | Biolegend #506329 |
| CD183 (CXCR3) | CXCR3-173 | BD Biosciences #740630 |
| CD279 (PD-1) | RMP1-30 | BD Biosciences #749306 |
| Granzyme-B | GB12 | Invitrogen #MHGB04 |
| T-bet | 4B10 | Invitrogen #25-5825-82 |
| CD3 | 145-2C11 | BD Biosciences #564661 |

Flow cytometry analysis was performed on BD FACS Symphony A5 SORP and analyzed with FlowJo software (Treestar).

### Tissue homogenate and viral titers

Tissues homogenates were prepared by homogenizing perfused lungs and brains using gentleMACS Octo Dissociator (Miltenyi BioTec, #130-096-427) in M tubes (#130-093-335) containing 1 ml of DMEM 0% FBS. Samples were homogenized for three times with program m_Lung_01_02 (34 seconds, 164 rpm). The homogenates were centrifuged at 3’500 rpm for 5 minutes at 4°C. The supernatant was collected and stored at −80°C for viral isolation and viral load detection. Viral titer was calculated by 50% tissue culture infectious dose (TCID50). Briefly, Vero E6 cells were seeded at a density of 1.5 × 10^4^ cells per well in flat-bottom 96-well tissue culture plates. The following day, 10-fold dilutions of the homogenized tissue were applied to confluent cells and incubated 1h at 37°C. Then, cells were washed with phosphate-buffered saline (PBS) and incubated for 72h at 37°C in DMEM 2% FBS. Cells were fixed with 4% paraformaldehyde for 30 min and stained with 0.05% (wt/vol) crystal violet in 20% ethanol.

### RNA extraction and qPCR

Tissues homogenates were prepared by homogenizing perfused lung, brain, olfactory bulb, and nasal turbinates using gentleMACS dissociator (Miltenyi BioTec, #130-096-427) with program RNA_02 in M tubes (#130-096-335) in 1 ml of Trizol (Invitrogen, #15596018). The homogenates were centrifuged at 2000 g for 1 min at 4°C and the supernatant was collected. RNA extraction was performed by combining phenol/guanidine-based lysis with silica membrane-based purification. Briefly, 100 μl of Chloroform were added to 500 μl of homogenized sample and total RNA was extracted using ReliaPrep™ RNA Tissue Miniprep column (Promega, Cat #Z6111). Total RNA was isolated according to the manufacturer’s instructions. qPCR was performed using TaqMan Fast virus 1 Step PCR Master Mix (Lifetechnologies #4444434), standard curve was drawn with 2019_nCOV_N Positive control (IDT#10006625), primer used are: 2019-nCoV_N1- Forward Primer (5’-GAC CCC AAA ATC AGC GAA AT-3’), 2019- nCoV_N1- Reverse Primer (5’-TCT GGT TAC TGC CAG TTG AAT CTG-3’) 2019- nCoV_N1-Probe (5’-FAM-ACC CCG CAT TAC GTT TGG TGG ACC-BHQ1-3’) (Centers for Disease Control and Prevention (CDC) Atlanta, GA 30333). All experiments were performed in duplicate.

### RNA-seq library preparation

Total RNA was obtained from homogenized lung tissues, as described above, for bulk RNA sequencing. Sequencing libraries were generated using Smart-seq2 method, as described (Picelli et al., 2014; Bénéchet et al., 2019). In brief, 4 ng of RNA were retrotranscribed and cDNA was amplified using 13 cycles and purified with AMPure XP beads (Beckman Coulter). Concentration was determined using Qubit 3.0 (Life Technologies) and the size distribution was assessed using Agilent 4200 TapeStation system. The following tagmentation reaction was performed starting from 0.5 ng of cDNA for 30 min at 55°C and the enrichment PCR was carried out using 12 cycles.

Libraries were then purified with AMPure XP beads, quantified using Qubit 3.0 and single-end sequenced (75 bp) on an Illumina NextSeq 500.

### RNA-seq data processing and analysis

Raw reads were aligned to mouse genome build GRCm38 using STAR aligner (Dobin et al., 2013). Gene counts were generated using featureCounts (part of the Subread package (Liao et al., 2019)), based on GENCODE gene annotation version M22. In order to discard genes highly expressed by one sample only, a filter of cpm *≥* 2 in at least two samples was added. Read counts were normalized with the Trimmed Mean of M-values (TMM) method (Robinson and Oshlack, 2010) using calcNormFactors function and then Voom (Law et al., 2014) was applied. Differentially Expressed Genes (DEGs) between AR- and IN-infected mice were identified by generating a linear model using LIMMA R package (Ritchie et al., 2015) (**Figure S6** and **Table S1**).

### Gene Set Enrichment Analysis and signatures visualization

Gene Set Enrichment Analysis (GSEA) was performed using the GseaPreranked Java tool (Subramanian et al., 2005) with pre-ranked Log2 fold changes between aerosol and intranasal samples in expressed genes. Three signatures described in (Winkler et al., 2020) were analyzed.

### Histochemistry

Mice were euthanized and perfused transcardially with PBS. One left lobe of the lung and a sagittal section of a hemisphere of the brain were fixed in zinc formalin and transferred into 70% ethanol 24h later. Tissues were then processed, embedded in paraffin, and automatically stained for SARS-CoV-2 (2019-nCoV) Nucleocapsid (SINO BIO, #40143-R019) or for fibrin (DAKO, #A0080) through LEICA BOND RX 1h room-temperature (RT) and developed with Bond Polymer Refine Detection (Leica, DS9800). For hematoxylin and eosin (H&E) staining, tissues were stained as previously described (Guidotti et al., 2015; Bénéchet et al., 2019). Bright-field images were acquired with an Aperio Scanscope System CS2 microscope and the ImageScope program (Leica Biosystem) following the manufacturer’s instructions. N-SARS-CoV-2 percentage of positive area (Figure S4 D, G) was determined by the QuPath (Quantitative Pathology & Bioimage 5 Analysis) software. Size of fibrin deposits (Figure 2 O) was determined by automatically building masks on fibrin-positive areas and calculating the covered surface that was expressed in μm^2^ with the QuPath software.

### Confocal Immunofluorescence Histology

Mice were euthanized and perfused transcardially with PBS. One left lobe of the lung was collected and fixed in 4% paraformaldehyde for 16h, then dehydrated in 30% sucrose prior to embedding in OCT freezing media (Killik Bio-Optica #05-9801). 20 μm sections were cut on a CM1520 cryostat (Leica) and adhered to Superfrost Plus slides (Thermo Scientific). Sections were permeabilized and blocked in PBS containing 0.3% Triton X-100 (Sigma-Aldrich) and 0,5% BSA followed by staining in PBS containing 0.1% Triton X-100 and 0,2% BSA. Slides were stained for SARS-CoV-2 nucleocapsid (GeneTex, polyclonal, #GTX135357), CD41 (Biolegend, Clone MWReg30, #133908) or TCRβ (Biolegend, Clone H57-597, #109218) for 1h at room temperature. Then, slides were stained with Alexa Fluor 568-conjugated goat anti-rabbit IgG (Life Technologies, #A-11011) for 2h at room temperature (for SARS-CoV-2 nucleocapsid staining). A hemisphere of the brain was fixed in 4% paraformaldehyde for 48h, then dehydrated in 30% sucrose for 24h. Prior to embedding in OCT freezing media, brain was soaked in a solution 1:1 of 30% sucrose and OCT for 1h in agitation. Brain was cut in 50 μm-thick sagittal sections and free-floating sections were rinsed in PBS/Azide 1%. Quenching was performed by shaking (300rpm) the sections for 10 min at room temperature in a PBS solution containing 1:100 methanol and 1:30 H202. The sections were then incubated 20 min with 0.3% Triton X-100 and 0.5% BSA for 1 h. Immunofluorescence staining was performed in PBS containing 0.1% Triton X-100 and 0.2% BSA over night (O/N) at 4°C and then stained with secondary antibody for 2h at room temperature. The following primary Abs were used for staining: anti-SARS-CoV-2 nucleocapsid (GeneTex, #GTX135357), anti-NeuN (Immulogical Sciences, #MAB377), anti-GFAP (Abcam, #ab4674), anti-Iba1 (Synaptic system, #234006), anti-CD68 (Abcam #ab53444), and anti-iNOS (Abcam #ab3523). The following secondary Abs were used for staining: Alexa Fluor 647-conjugated goat anti-chicken IgG (Thermo Fisher, #A-21), Alexa Fluor 568-conjugated goat anti-rabbit IgG (Thermo Fisher, #A-11011), Alexa Fluor 488-conjugated rat anti-mouse IgG (BD Bioscience, #553443). Lung and brain sections were washed twice for 5 min and stained with DAPI (Life technologies, #D1360) for 5 min at RT, then washed again and mounted for imaging with FluorSaveTM Reagent (Merck Millipore, #345789). Images were acquired on an SP5 or SP8 confocal microscope with 20x objective (Leica Microsystem). To minimize fluorophore spectral spillover, the Leica sequential laser excitation and detection modality was used.

### Statistical analyses and software

Detailed information concerning the statistical methods used is provided in the figure legends. Flow and imaging data were collected using FlowJo Version 10.5.3 (Treestar) and Imaris (Bitplane), respectively. Statistical analyses were performed with GraphPad Prism software version 8 (GraphPad). Immunohistochemical imaging quantifications were performed with QuPath (Quantitative Pathology & Bioimage 5 Analysis) software. *n* represents individual mice analyzed per experiment. Experiments were performed independently at least twice to control for experimental variation. Error bars indicate the standard error of the mean (SEM). We used Mann-Whitney U-tests to compare two groups with non-normally distributed continuous variables and Kruskal-Wallis non-parametric test to compare three or more unpaired groups. We used two-way ANOVA followed by Sidak’s multiple comparisons tests to analyze experiments with multiple groups and two independent variables. Kaplan-Meier curves were compared with the Log-rank (Mantel-Cox) test. Significance is indicated as follows: *p < 0.05; **p < 0.01; ***p < 0.001. Comparisons are not statistically significant unless indicated.

## Supplementary Figure Legends

**Figure S1.**
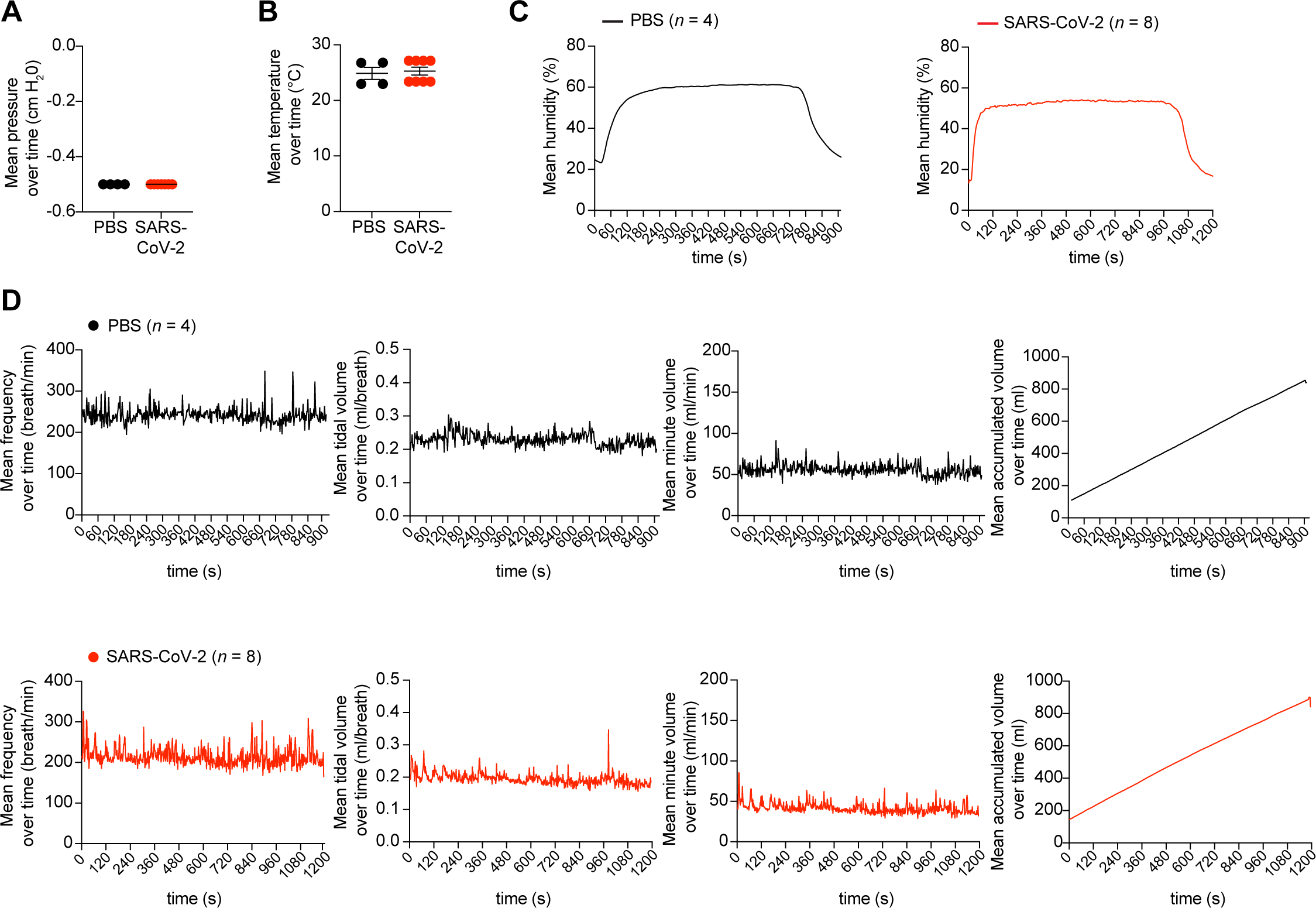
Recorded parameters in the nose-only inhalation tower system. **(A)** Mean of the pressure (cm H20) measured within the outer core of the nose-only inhalation tower during the time of PBS-exposure (*n* = 4, black dots) or SARS-CoV-2- exposure (*n* = 8, red dots) of K18-hACE2 mice. **(B)** Mean temperature (°C) over time at a chamber port exposed to PBS (*n* = 4, black dots) or SARS-CoV-2 (*n* = 8, red dots). **(C)** Mean humidity measured over time at a chamber port exposed to PBS (left panel, *n* = 4, black line) or SARS-CoV-2 (right panel, *n* = 8, red line). **(D)** Respiratory parameters analyzed during exposure through plethysmography associated to the Allay restrainer. Mean of breathing frequency, tidal volume, minute volume and accumulated inhaled volume measured during mouse exposure to PBS (upper panels, *n* = 4, black lines) or SARS-CoV-2 (lower panels, *n* = 8, red lines). Data are expressed as mean ± SEM (**A, B**) and as mean (**C-E**) and are representative of at least 2 independent experiments.

**Figure S2.**
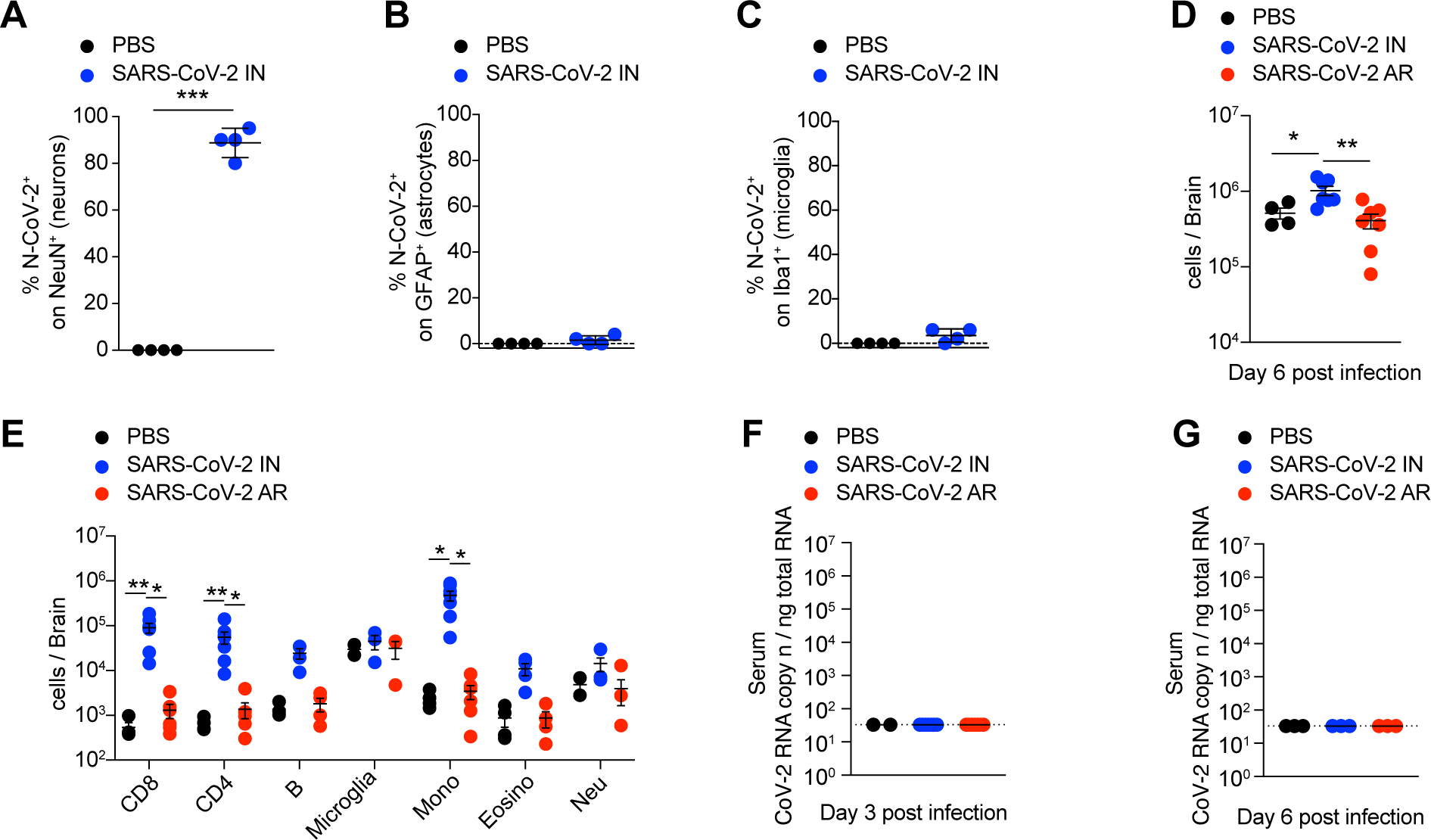
SARS-CoV-2 neuroinvasion occurs upon intranasal infection, but not upon aerosol exposure. **(A)** Quantification (corresponding to Figure 1L) of neurons infected with SARS-CoV-2 as percentage of N-CoV-2^+^ cells on total NeuN^+^ cells in the cerebral cortex of PBS-treated control mice (*n* = 4, black dots) and IN-infected mice (*n* = 4, blue dots) 6 days post infection. **(B)** Quantification (corresponding to Figure 1N) of astrocytes infected with SARS-CoV-2 as percentage of N-CoV-2^+^ cells on total GFAP^+^ cells in the cerebral cortex of PBS-treated control mice (*n* = 4, black dots) and IN- infected mice (*n* = 4, blue dots) 6 days post infection. **(C)** Quantification (corresponding to Figure 1O) of infected microglia as percentage of N-CoV-2^+^ cells on the total Iba1^+^ cells in the cerebral cortex of PBS-treated control mice (*n* = 4, black dots) and IN- infected mice (*n* = 4, blue dots) 6 days post infection. **(D, E)** Absolute number of total cells (**D**) and single cell population (**E**) recovered from brain homogenates of PBS- treated control mice (*n* = 4, black dots), IN- (*n* = 3-7, blue dots) and AR-infected mice (*n* = 3-7, red dots) analyzed 6 days post infection. CD8^+^ T cells (Live, CD45^+^, CD8^+^); CD4^+^ T cells (Live, CD45^+^, CD4^+^); B cells (Live, CD45^+^, CD8^-^, CD4^-^, B220^+^, CD19^+^); microglia (Live, CD45^int^, CD64^+^, F4/80^+^); Mono, monocytes (Live, CD45^hi^, CD8^-^, CD4^-^, Ly6g^-^, SiglecF^-^, CD11b^+^, Ly-6c^+^); Eosino, eosinophils (Live, CD45^hi^, CD8^-^, CD4^-^, Ly6g^-^, CD11b^+^, SiglecF^int^); Neu, neutrophils (Live, CD45^hi^, CD8^-^, CD4^-^, CD11b^+^, Ly6g^+^). **(F, G)** Quantification of SARS-CoV-2 RNA in the serum of IN- (*n* = 4, blue dots) and AR- (*n* = 4, red dots) infected mice as well as of PBS-treated control mice (*n* = 3, black dots) measured 3 days (**F**) and 6 days (**G**) post infection. RNA values are expressed as copy number per ng of total RNA and the limit of detection is indicated as a dotted line. Data are expressed as mean ± SEM. Data in (D-G) are pooled from 2 independent experiments per time point. * p-value < 0.05, ** p-value < 0.01, *** p-value < 0.001; Mann-Whitney U-test two-tailed (**A-C**); Kruskal-Wallis test (**D-G**).

**Figure S3.**
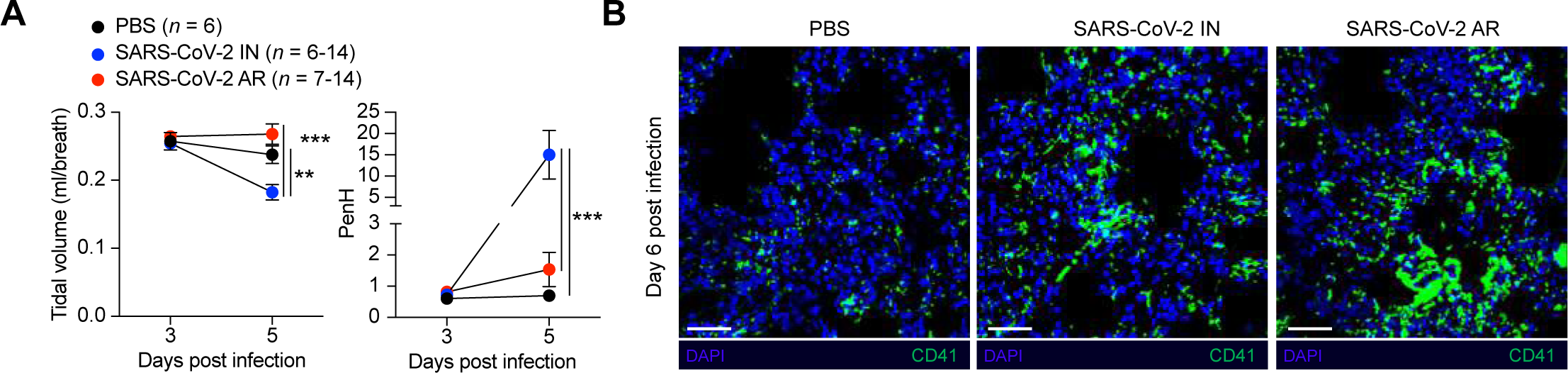
Breathing parameters and platelet aggregates in the lungs of SARS- CoV-2 infected mice. **(A)** Pulmonary function was assessed by whole-body plethysmography performed 3 and 5 days post IN- (*n* = 6-14, blue dots) and AR- infection (*n* = 7-14, red dots). As control, PBS-treated mice were evaluated (*n* = 6, black dots). Tidal volume (left) and PenH (right) parameters are shown. Calculated respiratory values were averaged over a 15 minute data collection period. **(B)** Representative confocal immunofluorescence micrographs of lung sections from PBS-treated control mice (left), IN- (middle) and AR-infected mice (right) 6 days post infection. CD41^+^ platelets are depicted in green; cell nuclei are depicted in blue. Scale bars, 30 μm. Data are expressed as mean ± SEM and are pooled from 2 independent experiments per time point. ** p-value < 0.01, *** p-value < 0.001; two-way ANOVA followed by Sidak’s multiple comparison test.

**Figure S4.**
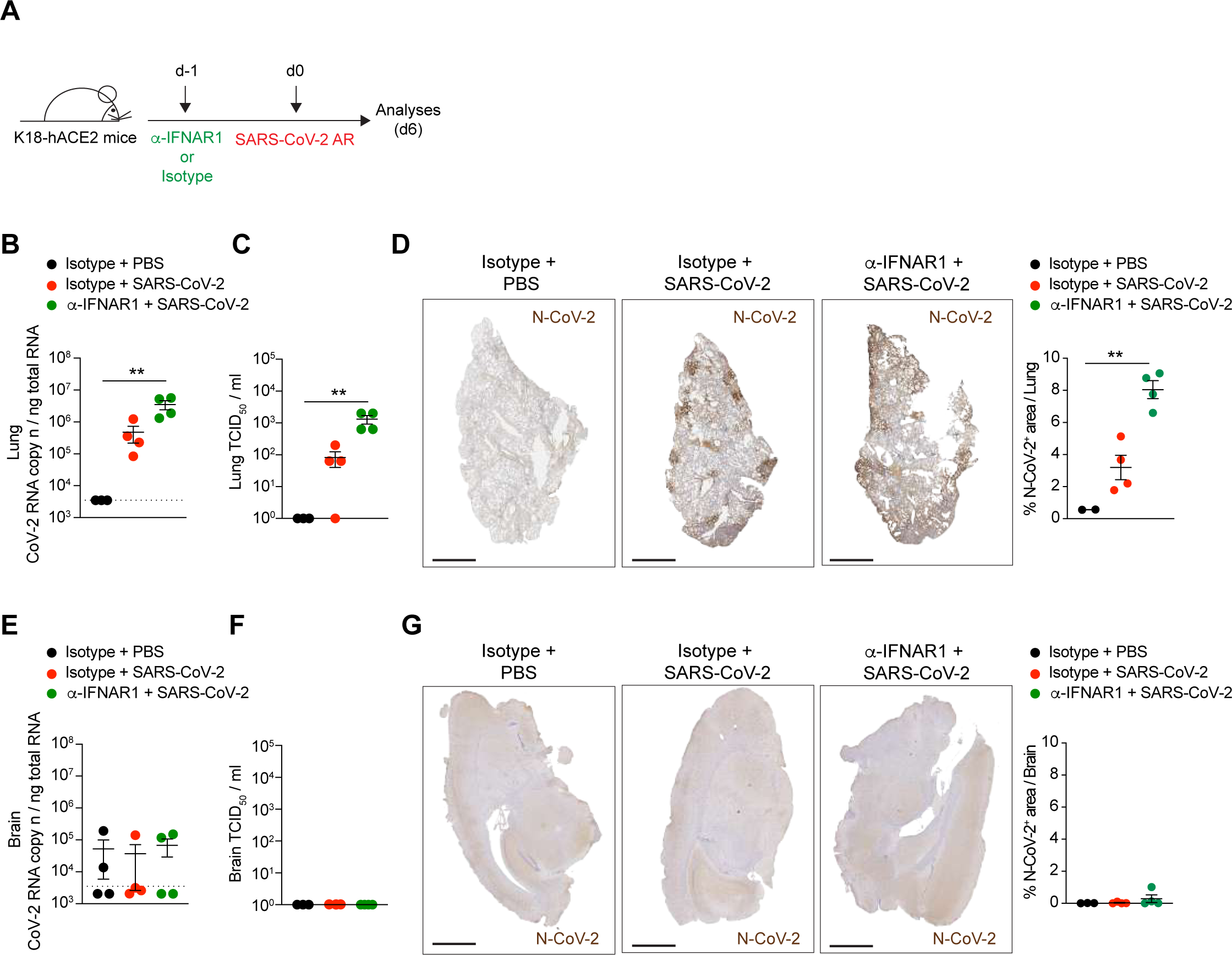
Aerosol exposure of K18-hACE2 transgenic mice to SARS-CoV-2 does not lead to fatal neuroinvasion even upon type I IFN receptor blockade. (A) Schematic representation of the experimental setup. K18h-ACE2 mice were treated with anti-IFNAR1 blocking antibody (or isotype control) 1 day before infection with a target dose of 1 x 10^5^ TCID50 of SARS-CoV-2 via aerosol exposure. Lungs and brains were collected and analyzed 6 days post infection. **(B, E)** Quantification of SARS-CoV-2 RNA in the lungs (**B**) and brains (**E**) of isotype control- (*n* = 4, red dots) and *α−*IFNAR1- (*n* = 4, green dots) treated mice as well as of PBS-treated control mice (*n* = 3, black dots) measured 6 days post infection. RNA values are expressed as copy number per ng of total RNA and the limit of detection is indicated as a dotted line. **(C, F)** Viral titers in the lungs (**C**) and brains (**F**) were determined 6 days after infection by median tissue culture infectious dose (TCID50). **(D, G)** Representative immunohistochemical micrographs of lung (**D**) and brain (**G**) sections from PBS-treated control mice (left), isotype control- (middle) and *α−*IFNAR1- (right) treated mice at 6 days post infection. N-CoV-2 positive cells are depicted in brown. Scale bars, 1 mm. Right panels, quantification of the percentage of N-CoV-2 positive area in the lung (top) and brain (bottom) of indicated mice; each dots represent one mouse. Data are expressed as mean ± SEM. ** p-value < 0.01; Kruskal-Wallis test (**B-G**).

**Figure S5.**
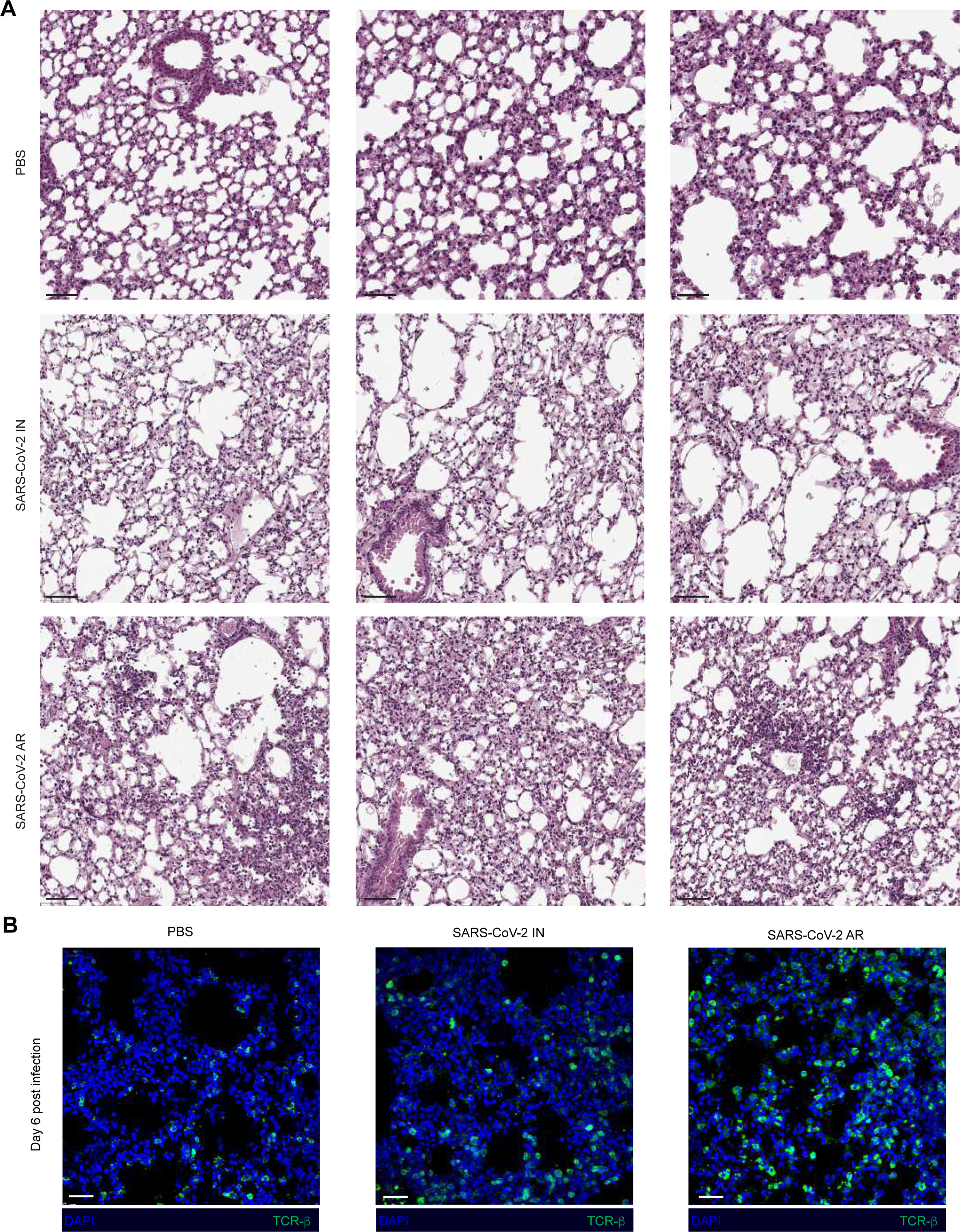
More severe lung pathology in AR-infected than in IN-infected mice. **(A)** Representative hematoxylin/eosin (H&E) micrographs of lung sections from PBS- treated control mice (top), IN- (middle) and AR-infected mice (bottom) 6 days post infection. Scale bars, 50 μm. **(B)** Representative confocal immunofluorescence micrographs of lung sections from PBS-treated control mice (left), IN- (middle) and AR- infected mice (right) 6 days post infection. TCR-b positive cells are depicted in green; cell nuclei are depicted in blue. Scale bars, 30 μm.

**Figure S6.**
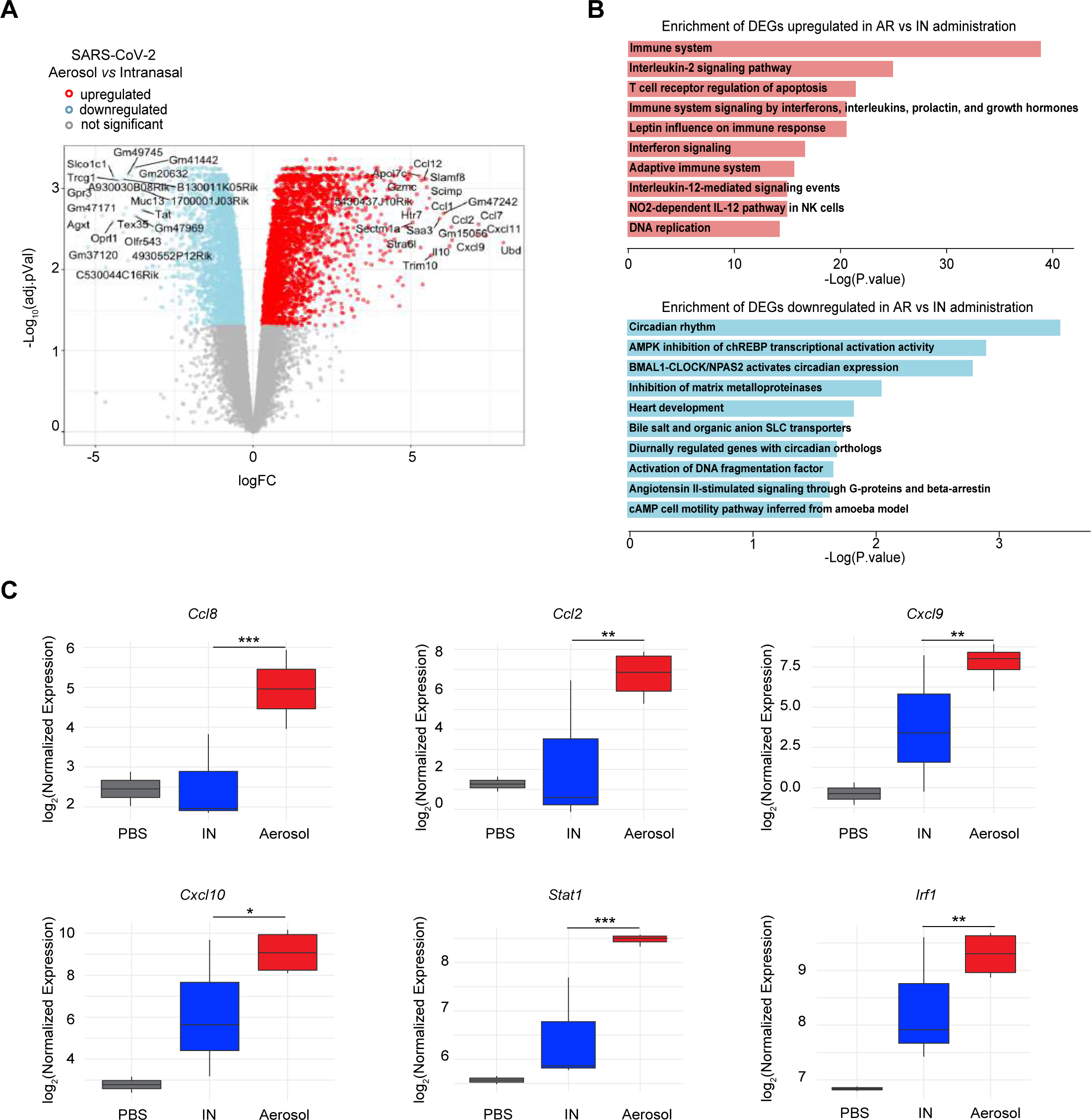
Transcriptional signature in the lungs of infected mice. **(A)** Volcano plot of RNA-seq results. The X-axis represents the Log2 Fold-Change of Differentially Expressed Genes (DEG) comparing AR- to IN-infected mice, the Y-axis the - Log10(FDR) Genes significantly upregulated in AR- relative to IN-infected mice (adjusted P value < 0.05) are colored in red and genes significantly downregulated are colored in blue. **(B)** Top ten pathways enriched by *p* values (resulting from a Fisher exact test) from BioPlanet 2019 database (Huang et al., 2019) for upregulated and downregulated genes (|logFC| > 1 and adjusted P value < 0.01). Enrichment tests were performed using the EnrichR web platform (https://maayanlab.cloud/Enrichr/). **(C)** Boxplot representing the expression level of the indicated genes in the lung of PBS-treated control mice (*n* = 2, gray boxes), IN- (*n* = 3, blue boxes) and AR-infected (*n* = 4, red boxes) mice 6 days post infection. Y-axis indicates the logarithmic normalized read counts. Comparison between IN-infection and AR-infection. * adjusted P value < 0.05, ** adjusted P value < 0.01, *** adjusted P value < 0.001. Adjusted P value corrected using Benjamini Hochberg correction method.

## Supplementary Table Legends

**Supplementary Table 1. *Differential Gene Expression between AR- and IN- infected mice.*** Results showed in Figure 3F-I and Figure S6. Genes expressed (cpm *≥* 2 in at least 2 samples) were processed using LIMMA (Ritchie et al., 2015) and significant differentially expressed genes (Adjusted P value < 0.05) are showed.

## References

1. Bates, J., C. Irvin, V. Brusasco, J. Drazen, J. Fredberg, S. Loring, D. Eidelman, M. Ludwig, P. Macklem, J. Martin, J. Milic-Emili, Z. Hantos, R. Hyatt, S. Lai-Fook, A. Leff, J. Solway, K. Lutchen, B. Suki, W. Mitzner, P. Paré, N. Pride, and P. Sly. 2004. The Use and Misuse of Penh in Animal Models of Lung Disease. Am J Resp Cell Mol. 31:373–374. doi:10.1165/ajrcmb.31.3.1.

2. Bénéchet, A.P., G.D. Simone, P.D. Lucia, F. Cilenti, G. Barbiera, N.L. Bert, V. Fumagalli, E. Lusito, F. Moalli, V. Bianchessi, F. Andreata, P. Zordan, E. Bono, L. Giustini, W.V. Bonilla, C. Bleriot, K. Kunasegaran, G. Gonzalez-Aseguinolaza, D.D. Pinschewer, P.T.F. Kennedy, L. Naldini, M. Kuka, F. Ginhoux, A. Cantore, A. Bertoletti, R. Ostuni, L.G. Guidotti, and M. Iannacone. 2019. Dynamics and genomic landscape of CD8+ T cells undergoing hepatic priming. Nature. 574:200–205. doi:10.1038/s41586-019-1620-6.

3. Blanco-Melo, D., B.E. Nilsson-Payant, W.-C. Liu, S. Uhl, D. Hoagland, R. Møller, T.X. Jordan, K. Oishi, M. Panis, D. Sachs, T.T. Wang, R.E. Schwartz, J.K. Lim, R.A. Albrecht, and B.R. tenOever. 2020. Imbalanced Host Response to SARS-CoV-2 Drives Development of COVID-19. Cell. 181:1036–1045.e9. doi:10.1016/j.cell.2020.04.026.

4. Bradley, B.T., H. Maioli, R. Johnston, I. Chaudhry, S.L. Fink, H. Xu, B. Najafian, G. Deutsch, J.M. Lacy, T. Williams, N. Yarid, and D.A. Marshall. 2020. Histopathology and ultrastructural findings of fatal COVID-19 infections in Washington State: a case series. Lancet. 396:320–332. doi:10.1016/s0140-6736(20)31305-2.

5. Chow, Y.-H., H. O’Brodovich, J. Plumb, Y. Wen, K.-J. Sohn, Z. Lu, F. Zhang, G.L. Lukacs, A.K. Tanswell, C.-C. Hui, M. Buchwald, and J. Hu. 1997. Development of an epithelium-specific expression cassette with human DNA regulatory elements for transgene expression in lung airways. Proc National Acad Sci. 94:14695–14700. doi:10.1073/pnas.94.26.14695.

6. Delorey, T.M., C.G.K. Ziegler, G. Heimberg, R. Normand, Y. Yang, Å. Segerstolpe, D. Abbondanza, S.J. Fleming, A. Subramanian, D.T. Montoro, K.A. Jagadeesh, K.K. Dey, P. Sen, M. Slyper, Y.H. Pita-Juárez, D. Phillips, J. Biermann, Z. Bloom-Ackermann, N. Barkas, A. Ganna, J. Gomez, J.C. Melms, I. Katsyv, E. Normandin, P. Naderi, Y.V. Popov, S.S. Raju, S. Niezen, L.T.-Y. Tsai, K.J. Siddle, M. Sud, V.M. Tran, S.K. Vellarikkal, Y. Wang, L. Amir-Zilberstein, D.S. Atri, J. Beechem, O.R. Brook, J. Chen, P. Divakar, P. Dorceus, J.M. Engreitz, A. Essene, D.M. Fitzgerald, R. Fropf, S. Gazal, J. Gould, J. Grzyb, T. Harvey, J. Hecht, T. Hether, J. Jané-Valbuena, M. Leney-Greene, H. Ma, C. McCabe, D.E. McLoughlin, E.M. Miller, C. Muus, M. Niemi, R. Padera, L. Pan, D. Pant, C. Pe’er, J. Pfiffner-Borges, C.J. Pinto, J. Plaisted, J. Reeves, M. Ross, M. Rudy, E.H. Rueckert, M. Siciliano, A. Sturm, E. Todres, A. Waghray, S. Warren, S. Zhang, D.R. Zollinger, L. Cosimi, R.M. Gupta, N. Hacohen, H. Hibshoosh, W. Hide, A.L. Price, J. Rajagopal, P.R. Tata, S. Riedel, G. Szabo, T.L. Tickle, P.T. Ellinor, D. Hung, P.C. Sabeti, R. Novak, R. Rogers, D.E. Ingber, Z.G. Jiang, D. Juric, M. Babadi, S.L. Farhi, et al. 2021. COVID-19 tissue atlases reveal SARS-CoV-2 pathology and cellular targets. Nature. 595:107–113. doi:10.1038/s41586-021-03570-8.

7. Dobin, A., C.A. Davis, F. Schlesinger, J. Drenkow, C. Zaleski, S. Jha, P. Batut, M. Chaisson, and T.R. Gingeras. 2013. STAR: ultrafast universal RNA-seq aligner. Bioinformatics. 29:15–21. doi:10.1093/bioinformatics/bts635.

8. Dolhnikoff, M., A.N. Duarte-Neto, R.A.A. Monteiro, L.F.F. Silva, E.P. Oliveira, P.H.N. Saldiva, T. Mauad, and E.M. Negri. 2020. Pathological evidence of pulmonary thrombotic phenomena in severe COVID-19. J Thromb Haemost. 18:1517–1519. doi:10.1111/jth.14844.

9. Fox, S.E., A. Akmatbekov, J.L. Harbert, G. Li, J.Q. Brown, and R.S.V. Heide. 2020. Pulmonary and cardiac pathology in African American patients with COVID-19: an autopsy series from New Orleans. Lancet Respir Medicine. 8:681–686. doi:10.1016/s2213-2600(20)30243-5.

10. Glatzel, M., C. Hagel, J. Matschke, J. Sperhake, N. Deigendesch, A. Tzankov, and S. Frank. 2021. Neuropathology associated with SARS-CoV-2 infection. Lancet. 397:276. doi:10.1016/s0140-6736(21)00098-2.

11. Goody, R.J., C.C. Hoyt, and K.L. Tyler. 2005. Reovirus infection of the CNS enhances iNOS expression in areas of virus-induced injury. Exp Neurol. 195:379–390. doi:10.1016/j.expneurol.2005.05.016.

12. Guidotti, L.G., D. Inverso, L. Sironi, P. Di Lucia, J. Fioravanti, L. Ganzer, A. Fiocchi, M. Vacca, R. Aiolfi, S. Sammicheli, M. Mainetti, T. Cataudella, A. Raimondi, G. Gonzalez-Aseguinolaza, U. Protzer, Z.M. Ruggeri, F.V. Chisari, M. Isogawa, G. Sitia, and M. Iannacone. 2015. Immunosurveillance of the Liver by Intravascular Effector CD8+ T Cells. Cell. 161:486–500. doi:10.1016/j.cell.2015.03.005.

13. Helms, J., S. Kremer, H. Merdji, R. Clere-Jehl, M. Schenck, C. Kummerlen, O. Collange, C. Boulay, S. Fafi-Kremer, M. Ohana, M. Anheim, and F. Meziani. 2020. Neurologic Features in Severe SARS-CoV-2 Infection. New Engl J Med. 382:2268–2270. doi:10.1056/nejmc2008597.

14. Huang, R., I. Grishagin, Y. Wang, T. Zhao, J. Greene, J.C. Obenauer, D. Ngan, D.-T. Nguyen, R. Guha, A. Jadhav, N. Southall, A. Simeonov, and C.P. Austin. 2019. The NCATS BioPlanet – An Integrated Platform for Exploring the Universe of Cellular Signaling Pathways for Toxicology, Systems Biology, and Chemical Genomics. Front Pharmacol. 10:445. doi:10.3389/fphar.2019.00445.

15. Iadecola, C., J. Anrather, and H. Kamel. 2020. Effects of COVID-19 on the nervous system. Cell. 183:16–27.e1. doi:10.1016/j.cell.2020.08.028.

16. Iannacone, M., G. Sitia, M. Isogawa, J.K. Whitmire, P. Marchese, F.V. Chisari, Z.M. Ruggeri, and L.G. Guidotti. 2008. Platelets prevent IFN-α/β-induced lethal hemorrhage promoting CTL-dependent clearance of lymphocytic choriomeningitis virus. Proc National Acad Sci. 105:629–634. doi:10.1073/pnas.0711200105.

17. Jiang, R.-D., M.-Q. Liu, Y. Chen, C. Shan, Y.-W. Zhou, X.-R. Shen, Q. Li, L. Zhang, Y. Zhu, H.-R. Si, Q. Wang, J. Min, X. Wang, W. Zhang, B. Li, H.-J. Zhang, R.S. Baric, P. Zhou, X.-L. Yang, and Z.-L. Shi. 2020. Pathogenesis of SARS-CoV-2 in transgenic mice expressing human angiotensin-converting enzyme 2. Cell. 182:50–58.e8. doi:10.1016/j.cell.2020.05.027.

18. Kantonen, J., S. Mahzabin, M.I. Mäyränpää, O. Tynninen, A. Paetau, N. Andersson, A. Sajantila, O. Vapalahti, O. Carpén, E. Kekäläinen, A. Kantele, and L. Myllykangas. 2020. Neuropathologic features of four autopsied COVID-19 patients. Brain Pathol. 30:1012–1016. doi:10.1111/bpa.12889.

19. Karki, R., B.R. Sharma, S. Tuladhar, E.P. Williams, L. Zalduondo, P. Samir, M. Zheng, B. Sundaram, B. Banoth, R.K.S. Malireddi, P. Schreiner, G. Neale, P. Vogel, R. Webby, C.B. Jonsson, and T.-D. Kanneganti. 2021. Synergism of TNF-α and IFN-γ Triggers Inflammatory Cell Death, Tissue Damage, and Mortality in SARS-CoV-2 Infection and Cytokine Shock Syndromes. Cell. 184:149–168.e17. doi:10.1016/j.cell.2020.11.025.

20. Kumari, P., H.A. Rothan, J.P. Natekar, S. Stone, H. Pathak, P.G. Strate, K. Arora, M.A. Brinton, and M. Kumar. 2021. Neuroinvasion and Encephalitis Following Intranasal Inoculation of SARS-CoV-2 in K18-hACE2 Mice. Viruses. 13:132. doi:10.3390/v13010132.

21. Law, C.W., Y. Chen, W. Shi, and G.K. Smyth. 2014. voom: precision weights unlock linear model analysis tools for RNA-seq read counts. Genome Biol. 15:R29. doi:10.1186/gb-2014-15-2-r29.

22. Lax, S.F., K. Skok, P. Zechner, H.H. Kessler, N. Kaufmann, C. Koelblinger, K. Vander, U. Bargfrieder, and M. Trauner. 2020. Pulmonary Arterial Thrombosis in COVID-19 With Fatal Outcome: Results From a Prospective, Single-Center, Clinicopathologic Case Series. Ann Intern Med. 173:350–361. doi:10.7326/m20-2566.

23. Leist, S.R., K.H. Dinnon, A. Schäfer, L.V. Tse, K. Okuda, Y.J. Hou, A. West, C.E. Edwards, W. Sanders, E.J. Fritch, K.L. Gully, T. Scobey, A.J. Brown, T.P. Sheahan, N.J. Moorman, R.C. Boucher, L.E. Gralinski, S.A. Montgomery, and R.S. Baric. 2020. A Mouse-adapted SARS-CoV-2 induces Acute Lung Injury (ALI) and mortality in Standard Laboratory Mice. Cell. doi:10.1016/j.cell.2020.09.050.

24. Liao, Y., G.K. Smyth, and W. Shi. 2019. The R package Rsubread is easier, faster, cheaper and better for alignment and quantification of RNA sequencing reads. Nucleic Acids Res. 47:gkz114-. doi:10.1093/nar/gkz114.

25. Lomask, M. 2006. Further exploration of the Penh parameter. Exp Toxicol Pathol. 57:13–20. doi:10.1016/j.etp.2006.02.014.

26. Lundblad, L.K.A., C.G. Irvin, Z. Hantos, P. Sly, W. Mitzner, and J.H.T. Bates. 2007. Penh is not a measure of airway resistance! Eur Respir J. 30:805–805. doi:10.1183/09031936.00091307.

27. Manne, B.K., F. Denorme, E.A. Middleton, I. Portier, J.W. Rowley, C. Stubben, A.C. Petrey, N.D. Tolley, L. Guo, M. Cody, A.S. Weyrich, C.C. Yost, M.T. Rondina, and R.A. Campbell. 2020. Platelet gene expression and function in patients with COVID-19. Blood. 136:1317– 1329. doi:10.1182/blood.2020007214.

28. Mast, A.E., A.S. Wolberg, D. Gailani, M.R. Garvin, C. Alvarez, J.I. Miller, B. Aronow, and D. Jacobson. 2021. SARS-CoV-2 suppresses anticoagulant and fibrinolytic gene expression in the lung. Elife. 10:e64330. doi:10.7554/elife.64330.

29. Matschke, J., M. Lütgehetmann, C. Hagel, J.P. Sperhake, A.S. Schröder, C. Edler, H. Mushumba, A. Fitzek, L. Allweiss, M. Dandri, M. Dottermusch, A. Heinemann, S. Pfefferle, M. Schwabenland, D.S. Magruder, S. Bonn, M. Prinz, C. Gerloff, K. Püschel, S. Krasemann, M. Aepfelbacher, and M. Glatzel. 2020. Neuropathology of patients with COVID-19 in Germany: a post-mortem case series. Lancet Neurology. 19:919–929. doi:10.1016/s1474-4422(20)30308-2.

30. McCray, P.B., L. Pewe, C. Wohlford-Lenane, M. Hickey, L. Manzel, L. Shi, J. Netland, H.P. Jia, C. Halabi, C.D. Sigmund, D.K. Meyerholz, P. Kirby, D.C. Look, and S. Perlman. 2007. Lethal Infection of K18-hACE2 Mice Infected with Severe Acute Respiratory Syndrome Coronavirus. J Virol. 81:813–821. doi:10.1128/jvi.02012-06.

31. Meinhardt, J., J. Radke, C. Dittmayer, J. Franz, C. Thomas, R. Mothes, M. Laue, J. Schneider, S. Brünink, S. Greuel, M. Lehmann, O. Hassan, T. Aschman, E. Schumann, R.L. Chua, C. Conrad, R. Eils, W. Stenzel, M. Windgassen, L. Rößler, H.-H. Goebel, H.R. Gelderblom, H. Martin, A. Nitsche, W.J. Schulz-Schaeffer, S. Hakroush, M.S. Winkler, B. Tampe, F. Scheibe, P. Körtvélyessy, D. Reinhold, B. Siegmund, A.A. Kühl, S. Elezkurtaj, D. Horst, L. Oesterhelweg, M. Tsokos, B. Ingold-Heppner, C. Stadelmann, C. Drosten, V.M. Corman, H. Radbruch, and F.L. Heppner. 2021. Olfactory transmucosal SARS-CoV-2 invasion as a port of central nervous system entry in individuals with COVID-19. Nat Neurosci. 24:168–175. doi:10.1038/s41593-020-00758-5.

32. Melo, G.D. de, F. Lazarini, S. Levallois, C. Hautefort, V. Michel, F. Larrous, B. Verillaud, C. Aparicio, S. Wagner, G. Gheusi, L. Kergoat, E. Kornobis, F. Donati, T. Cokelaer, R. Hervochon, Y. Madec, E. Roze, D. Salmon, H. Bourhy, M. Lecuit, and P.-M. Lledo. 2021. COVID-19–related anosmia is associated with viral persistence and inflammation in human olfactory epithelium and brain infection in hamsters. Sci Transl Med. 13:eabf8396. doi:10.1126/scitranslmed.abf8396.

33. Menachery, V.D., L.E. Gralinski, R.S. Baric, and M.T. Ferris. 2015. New Metrics for Evaluating Viral Respiratory Pathogenesis. Plos One. 10:e0131451. doi:10.1371/journal.pone.0131451.

34. Muñoz-Fontela, C., W.E. Dowling, S.G.P. Funnell, P.-S. Gsell, A.X. Riveros-Balta, R.A. Albrecht, H. Andersen, R.S. Baric, M.W. Carroll, M. Cavaleri, C. Qin, I. Crozier, K. Dallmeier, L. de Waal, E. de Wit, L. Delang, E. Dohm, W.P. Duprex, D. Falzarano, C.L. Finch, M.B. Frieman, B.S. Graham, L.E. Gralinski, K. Guilfoyle, B.L. Haagmans, G.A. Hamilton, A.L. Hartman, S. Herfst, S.J.F. Kaptein, W.B. Klimstra, I. Knezevic, P.R. Krause, J.H. Kuhn, R.L. Grand, M.G. Lewis, W.-C. Liu, P. Maisonnasse, A.K. McElroy, V. Munster, N. Oreshkova, A.L. Rasmussen, J. Rocha-Pereira, B. Rockx, E. Rodríguez, T.F. Rogers, F.J. Salguero, M. Schotsaert, K.J. Stittelaar, H.J. Thibaut, C.-T. Tseng, J. Vergara-Alert, M. Beer, T. Brasel, J.F.W. Chan, A. García-Sastre, J. Neyts, S. Perlman, D.S. Reed, J.A. Richt, C.J. Roy, J. Segalés, S.S. Vasan, A.M. Henao-Restrepo, and D.H. Barouch. 2020. Animal models for COVID-19. Nature. 586:509–515. doi:10.1038/s41586-020-2787-6.

35. Picelli, S., O.R. Faridani, Å.K. Björklund, G. Winberg, S. Sagasser, and R. Sandberg. 2014. Full-length RNA-seq from single cells using Smart-seq2. Nat Protoc. 9:171–181. doi:10.1038/nprot.2014.006.

36. Puelles, V.G., M. Lütgehetmann, M.T. Lindenmeyer, J.P. Sperhake, M.N. Wong, L. Allweiss, S. Chilla, A. Heinemann, N. Wanner, S. Liu, F. Braun, S. Lu, S. Pfefferle, A.S. Schröder, C. Edler, O. Gross, M. Glatzel, D. Wichmann, T. Wiech, S. Kluge, K. Pueschel, M. Aepfelbacher, and T.B. Huber. 2020. Multiorgan and Renal Tropism of SARS-CoV-2. New Engl J Med. 383:590–592. doi:10.1056/nejmc2011400.

37. Qiu, C., C. Cui, C. Hautefort, A. Haehner, J. Zhao, Q. Yao, H. Zeng, E.J. Nisenbaum, L. Liu, Y. Zhao, D. Zhang, C.G. Levine, I. Cejas, Q. Dai, M. Zeng, P. Herman, C. Jourdaine, K. de With, J. Draf, B. Chen, D.T. Jayaweera, J.C. Denneny, R. Casiano, H. Yu, A.A. Eshraghi, T. Hummel, X. Liu, Y. Shu, and H. Lu. 2020. Olfactory and Gustatory Dysfunction as an Early Identifier of COVID-19 in Adults and Children: An International Multicenter Study. Otolaryngology Head Neck Surg. 163:714–721. doi:10.1177/0194599820934376.

38. Ramani, A., L. Müller, P.N. Ostermann, E. Gabriel, P. Abida-Islam, A. Müller-Schiffmann, A. Mariappan, O. Goureau, H. Gruell, A. Walker, M. Andrée, S. Hauka, T. Houwaart, A. Dilthey, K. Wohlgemuth, H. Omran, F. Klein, D. Wieczorek, O. Adams, J. Timm, C. Korth, H. Schaal, and J. Gopalakrishnan. 2020. SARS-CoV-2 targets neurons of 3D human brain organoids. Embo J. 39:e2020106230. doi:10.15252/embj.2020106230.

39. Reichard, R.R., K.B. Kashani, N.A. Boire, E. Constantopoulos, Y. Guo, and C.F. Lucchinetti. 2020. Neuropathology of COVID-19: a spectrum of vascular and acute disseminated encephalomyelitis (ADEM)-like pathology. Acta Neuropathol. 140:1–6. doi:10.1007/s00401-020-02166-2.

40. Ritchie, M.E., B. Phipson, D. Wu, Y. Hu, C.W. Law, W. Shi, and G.K. Smyth. 2015. limma powers differential expression analyses for RNA-sequencing and microarray studies. Nucleic Acids Res. 43:e47–e47. doi:10.1093/nar/gkv007.

41. Robinson, M.D., and A. Oshlack. 2010. A scaling normalization method for differential expression analysis of RNA-seq data. Genome Biol. 11:R25. doi:10.1186/gb-2010-11-3-r25.

42. Serrano, G.E., J.E. Walker, R. Arce, M.J. Glass, D. Vargas, L.I. Sue, A.J. Intorcia, C.M. Nelson, J. Oliver, J. Papa, A. Russell, K.E. Suszczewicz, C.I. Borja, C. Belden, D. Goldfarb, D. Shprecher, A. Atri, C.H. Adler, H.A. Shill, E. Driver-Dunckley, S.H. Mehta, B. Readhead, M.J. Huentelman, J.L. Peters, E. Alevritis, C. Bimi, J.P. Mizgerd, E.M. Reiman, T.J. Montine, M. Desforges, J.L. Zehnder, M.K. Sahoo, H. Zhang, D. Solis, B.A. Pinsky, M. Deture, D.W. Dickson, and T.G. Beach. 2021. Mapping of SARS-CoV-2 Brain Invasion and Histopathology in COVID-19 Disease. Medrxiv. 2021.02.15.21251511. doi:10.1101/2021.02.15.21251511.

43. Song, E., C. Zhang, B. Israelow, A. Lu-Culligan, A.V. Prado, S. Skriabine, P. Lu, O.-E. Weizman, F. Liu, Y. Dai, K. Szigeti-Buck, Y. Yasumoto, G. Wang, C. Castaldi, J. Heltke, E. Ng, J. Wheeler, M.M. Alfajaro, E. Levavasseur, B. Fontes, N.G. Ravindra, D.V. Dijk, S. Mane, M. Gunel, A. Ring, S.A.J. Kazmi, K. Zhang, C.B. Wilen, T.L. Horvath, I. Plu, S. Haik, J.-L. Thomas, A. Louvi, S.F. Farhadian, A. Huttner, D. Seilhean, N. Renier, K. Bilguvar, and A. Iwasaki. 2021. Neuroinvasion of SARS-CoV-2 in human and mouse brain. J Exp Med. 218:e20202135. doi:10.1084/jem.20202135.

44. Subramanian, A., P. Tamayo, V.K. Mootha, S. Mukherjee, B.L. Ebert, M.A. Gillette, A. Paulovich, S.L. Pomeroy, T.R. Golub, E.S. Lander, and J.P. Mesirov. 2005. Gene set enrichment analysis: A knowledge-based approach for interpreting genome-wide expression profiles. P Natl Acad Sci Usa. 102:15545–15550. doi:10.1073/pnas.0506580102.

45. Sun, S.-H., Q. Chen, H.-J. Gu, G. Yang, Y.-X. Wang, X.-Y. Huang, S.-S. Liu, N.-N. Zhang, X.-F. Li, R. Xiong, Y. Guo, Y.-Q. Deng, W.-J. Huang, Q. Liu, Q.-M. Liu, Y.-L. Shen, Y. Zhou, X. Yang, T.-Y. Zhao, C.-F. Fan, Y.-S. Zhou, C.-F. Qin, and Y.-C. Wang. 2020. A mouse model of SARS-CoV-2 infection and pathogenesis. Cell Host Microbe. 28:124–133.e4. doi:10.1016/j.chom.2020.05.020.

46. Winkler, E.S., A.L. Bailey, N.M. Kafai, S. Nair, B.T. McCune, J. Yu, J.M. Fox, R.E. Chen, J.T. Earnest, S.P. Keeler, J.H. Ritter, L.-I. Kang, S. Dort, A. Robichaud, R. Head, M.J. Holtzman, and M.S. Diamond. 2020. SARS-CoV-2 infection of human ACE2-transgenic mice causes severe lung inflammation and impaired function. Nat Immunol. 1–9. doi:10.1038/s41590-020-0778-2.

47. Wu, F., S. Zhao, B. Yu, Y.-M. Chen, W. Wang, Z.-G. Song, Y. Hu, Z.-W. Tao, J.-H. Tian, Y.-Y. Pei, M.-L. Yuan, Y.-L. Zhang, F.-H. Dai, Y. Liu, Q.-M. Wang, J.-J. Zheng, L. Xu, E.C. Holmes, and Y.-Z. Zhang. 2020. A new coronavirus associated with human respiratory disease in China. Nature. 579:265–269. doi:10.1038/s41586-020-2008-3.

48. Xydakis, M.S., P. Dehgani-Mobaraki, E.H. Holbrook, U.W. Geisthoff, C. Bauer, C. Hautefort, P. Herman, G.T. Manley, D.M. Lyon, and C. Hopkins. 2020. Smell and taste dysfunction in patients with COVID-19. Lancet Infect Dis. 20:1015–1016. doi:10.1016/s1473-3099(20)30293-0.

49. Zheng, J., L.-Y.R. Wong, K. Li, A.K. Verma, M.E. Ortiz, C. Wohlford-Lenane, M.R. Leidinger, C.M. Knudson, D.K. Meyerholz, P.B. McCray, and S. Perlman. 2021. COVID-19 treatments and pathogenesis including anosmia in K18-hACE2 mice. Nature. 589:603–607. doi:10.1038/s41586-020-2943-z.

50. Zhou, L., S.K. Ayeh, V. Chidambaram, and P.C. Karakousis. 2021. Modes of transmission of SARS-CoV-2 and evidence for preventive behavioral interventions. Bmc Infect Dis. 21:496. doi:10.1186/s12879-021-06222-4.

51. Zhou, P., X.-L. Yang, X.-G. Wang, B. Hu, L. Zhang, W. Zhang, H.-R. Si, Y. Zhu, B. Li, C.-L. Huang, H.-D. Chen, J. Chen, Y. Luo, H. Guo, R.-D. Jiang, M.-Q. Liu, Y. Chen, X.-R. Shen, X. Wang, X.-S. Zheng, K. Zhao, Q.-J. Chen, F. Deng, L.-L. Liu, B. Yan, F.-X. Zhan, Y.-Y. Wang, G.-F. Xiao, and Z.-L. Shi. 2020. A pneumonia outbreak associated with a new coronavirus of probable bat origin. Nature. 579:270–273. doi:10.1038/s41586-020-2012-7.

52. Zhuang, Z., X. Lai, J. Sun, Z. Chen, Z. Zhang, J. Dai, D. Liu, Y. Li, F. Li, Y. Wang, A. Zhu, J. Wang, W. Yang, J. Huang, X. Li, L. Hu, L. Wen, J. Zhuo, Y. Zhang, D. Chen, S. Li, S. Huang, Y. Shi, K. Zheng, N. Zhong, J. Zhao, D. Zhou, and J. Zhao. 2021. Mapping and role of T cell response in SARS-CoV-2–infected mice. J Exp Med. 218:e20202187. doi:10.1084/jem.20202187.

